# VASP mediated actin dynamics activate and recruit a filopodia myosin

**DOI:** 10.1101/2021.03.16.435667

**Authors:** Ashley L. Arthur, Amy Crawford, Anne Houdusse, Margaret A. Titus

**Affiliations:** Department of Genetics, Cell Biology, and Development, University of Minnesota, Minneapolis, MN 55455; Structural Motility, Institut Curie, Paris Université Sciences et Lettres, Sorbonne Université, CNRS UMR144, 75005 Paris

**Keywords:** Filopodia, actin dynamics, VASP, MyTH4-FERM myosin

## Abstract

Filopodia are thin, actin-based structures that cells use to interact with their environments. Filopodia initiation requires a suite of conserved proteins but the mechanism remains poorly understood. The actin polymerase VASP and a MyTH-FERM (MF) myosin, DdMyo7 in amoeba, are essential for filopodia initiation. DdMyo7 is localized to dynamic regions of the actin-rich cortex. Analysis of VASP mutants and treatment of cells with anti-actin drugs shows that myosin recruitment and activation in *Dictyostelium* requires localized VASP-dependent actin polymerization. Targeting of DdMyo7 to the cortex alone is not sufficient for filopodia initiation; VASP activity is also required. The actin regulator locally produces a cortical actin network that activates myosin and together they shape the actin network to promote extension of parallel bundles of actin during filopodia formation. This work reveals how filopodia initiation requires close collaboration between an actin binding protein, the state of the actin cytoskeleton and MF myosin activity.

## Introduction

The efficient and directed migration of cells depends on their ability to detect and respond to chemical signals and physical cues in the environment. Filopodia are dynamic, thin membrane projections supported by a parallel bundle of actin filaments. They detect extracellular cues and play roles in processes such as neuronal growth cone guidance, durotaxis, cell-cell junction formation during development and metastasis (Heckman and Plummer, 2013; Arjonen et al., 2011; Cao et al., 2014; Shibue et al., 2012; Gallop, 2020). Although most intensely studied in animal cells, filopodia are ubiquitous in moving cells and have been observed in various Rhizaria, including predatory vampire amoebae, Discoba, Apusoza, Amoeboza and Holozoa (Sebé-Pedrós et al., 2013; Cavalier-Smith and Chao, 2003; Hess et al., 2012; Hanousková et al., 2019; Yabuki et al., 2013). Filopodia formation is orchestrated by a conserved core set of proteins that drive the formation and extension of actin bundles. These include a Rho family GTPase (Rac1, Cdc42), an actin polymerase (VASP or formin), an actin cross-linker and a MyTH4-FERM Myosin (MF; **my**osin **t**ail **h**omology 4, band **4**.1, **e**zrin, **r**adixin, **m**oesin) (Mattila et al., 2007; Sebé-Pedrós et al., 2013; Nobes and Hall, 1995; Tuxworth et al., 2001; Faix et al., 2009). MF myosin motors regulate the formation of filopodia and other parallel actin based structures (Weck et al., 2017). *Dictyostelium* amoebae null for DdMyo7 do not produce filopodia or any filopodia-like protrusions, and expression of Myo10 in various mammalian cell types induces filopodia formation, implicating these myosins in filopodia initiation (Bohil et al., 2006; Tuxworth et al., 2001; Sousa and Cheney, 2005). These MF myosins are strikingly localized to filopodia tips, yet their function during filopodia initiation remains poorly defined.

Filopodia protrude from cells as slender projections filled with parallel actin filaments. Actin polymerization aided by VASP or formin facilitates the formation of a critical bundle size of 15- 20 filaments which overcomes membrane tension and extends outwards as the actin filaments are cross-linked together (Mattila and Lappalainen, 2008; Mogilner and Rubinstein, 2005). Two models have been proposed for how these parallel actin bundles are formed. In one case the branched actin network is reorganized into a parallel array by the polymerase and bundler VASP (Svitkina et al., 2003; Yang and Svitkina, 2011). In an alternative model, a linear actin polymerase such as formin nucleates new filaments that are rapidly bundled together and grow perpendicular to the plasma membrane (Faix and Rottner, 2006). MF myosins are thought to act during initiation by cross-linking actin filaments, perhaps zipping them together as the motors walk up the filament towards the cortex (Ropars et al., 2016). Support for this model comes from the observation that forced dimers of motors can induce filopodia or filopodia-like protrusions in cells (Tokuo et al., 2007; Arthur et al., 2019; Masters and Buss, 2017; Liu et al., 2021). Myo10 has also been implicated in elongation by transporting VASP towards the tip of the growing filopodium to promote continued growth (Tokuo and Ikebe, 2004).

The mechanism by which MF myosins are recruited to filopodial initiation sites is not well-understood. These myosins are regulated by head-tail autoinhibition, with the myosin folded into a compact conformation whereby binding of the C-terminal MF domain to the motor domain inhibits its activity. Opening up of the myosin followed by dimerization is required for activation of the myosin (Umeki et al., 2011; Sakai et al., 2011; Yang et al., 2006; Arthur et al., 2019). Partner binding mediated by their MF domains can typically stabilize the open, activated form of these myosins as well as promote dimerization (Sakai et al., 2011; Arthur et al., 2019; Liu et al., 2021). The MyTH4-FERM (MF) domains in these myosins can indeed mediate interaction with partner proteins such as microtubules (Weber et al., 2004; Toyoshima and Nishida, 2007; Planelles-Herrero et al., 2016) and the cytoplasmic tails of adhesion and signaling receptors (Hirao et al., 1996; Hamada, 2000; Zhang et al., 2004; Zhu et al., 2007). Myo10 has a unique tail among MF myosins with PH domains that bind to PIP3 rich membranes, which facilitates autoinhibition release and subsequent dimerization via a coiled-coil domain (Umeki et al., 2011; Lu et al., 2012; Ropars et al., 2016). DdMyo7, like mammalian microvilli and stereocilia myosins Myo7 and Myo15, lacks PH domains and it is not clear if activation is regulated by the concentration of some partners, or additional cellular signals.

MF myosin and the actin regulators VASP and formin have been observed to coalesce into punctae during the initiation step that precedes extension (Young et al., 2018; Arthur et al., 2019; Cheng and Mullins, 2020). The mechanism by which Myo7s are targeted to such sites is unknown, but the first step of this process is likely to be regulated via relief of autoinhibition. Characterization of filopodia formation in *Dictyostelium* allows for a systematic examination of how DdMyo7 is recruited to the cell cortex through identification of the essential features of this myosin. The mechanism of DdMyo7 recruitment to the cortex and its potential functional relationship with the actin polymerase and bundler VASP were investigated to gain new insights into motor activation and the early steps of filopodia formation.

## Results

### The DdMyo7 motor restricts cortical localization of the tail

Disruption of the *Dictyostelium myo7* gene encoding DdMyo7 results in a significant defect in filopodia formation that is rescued by expression of GFP-DdMyo7 (Figure 1A, Supplemental Figure 1A and (Tuxworth et al., 2001; Petersen et al., 2016)). Most strikingly localized to filopodia tips, DdMyo7 is also in the cytosol and localized to the leading edge of cells (Figure 1B, Supplemental Figure 1A). In the course of characterizing functionally important regions of DdMyo7, it was observed that the tail domain (aa 809 - end) is localized all around the cell periphery in contrast to the full-length myosin which is often restricted at one edge of the cell cortex, (Figure 1B; (Petersen et al., 2016; Arthur et al., 2019)). This was unexpected as the myosin tail region is largely regarded as playing a key, even determining, role in targeting myosins. DdMyo7-mCherry and GFP- tail were co-expressed in *Dictyostelium* cells and a line scan through the cell showed that while both are present at a region of the cell that is extending outwards and producing filopodia (i.e. the leading edge), the tail is also strongly enriched in the cell rear (Figure 1C, line from Figure 1B). The extent of co-localization of the full-length myosin with the tail domain was assessed using cytofluorograms. This method quantifies co-localization of two proteins by comparing the intensity of the two fluorescent signals on a pixel-wise basis for the entire image with *r* representing the correlation coefficient (Bolte and Cordelières, 2006). Quantification revealed significantly less correlation between DdMyo7 and the tail (r = 0.63) than is seen for the two full-length myosins (r = 0.86). Due to the transient polarity of vegetative cells, a second measure of distribution of DdMyo7 on the cortex was carried out. To quantify the localized enrichment of DdMyo7 without any bias for manually identifying the leading edge, the intensity all around the cortex of cells was measured using a high throughput automated image analysis FIJI macro, *Seven* (Petersen et al., 2016) (Figure 1E). Actin is typically enriched at the leading edge with high intensity at one edge and lower intensity elsewhere (Figure 1F, G). The standard deviation of intensities values around the periphery was calculated for each cell (as exemplified in Figure 1H). This analysis shows that actin (RFP-Lifeact) and GFP-DdMyo7 are localized asymmetrically with a high standard deviation of cortical intensity (Figure 1F, G, I). In contrast, the tail appears uniformly localized around the periphery (Figure 1F, G) and has a lower average SD for the cortical intensity around the cell periphery (Figure 1I). Interestingly, a tail-less forced dimer of the motor (motor-FD; Figure 1A) that is capable of inducing filopodial protrusions (Arthur et al., 2019) is also localized asymmetrically with a high standard deviation of intensity (Figure 1F, I). Together, these data establish that restricted localization of full-length DdMyo7 is dependent on its motor domain.

**Figure 1.**
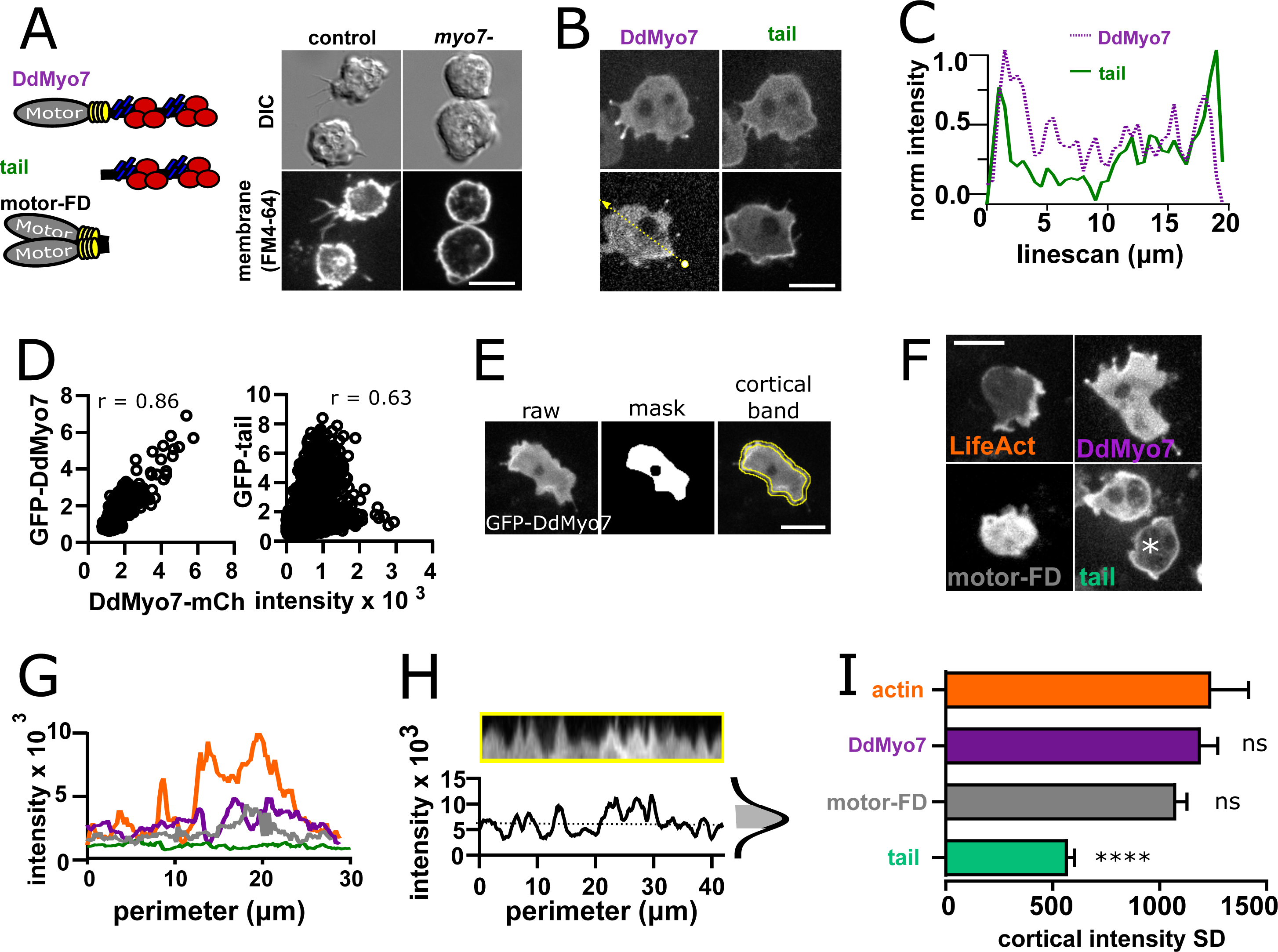
DdMyo7 has a distinct cortical localization from its tail domain. **A.** (left) Schematic of DdMyo7 illustrating its motor domain (grey), 4 IQ domains (yellow) and tandem MyTH4- FERM domains (blue-MyTH4, red-FERM) in the tail, the tail fragment, and a motor forced dimer (motor-FD); (right) *Dictyostelium* control, or *myo7* null cells visualized with DIC and the membrane dye FM4-64 showing DdMyo7 is critical for filopodia formation. **B.** Confocal images showing localization of DdMyo7-mCherry at the cortex and in filopodia tips, and GFP-tail fragment localized around cortex. **C.** Line intensity profile along the line shown in panel B. **D.** Cytofluorograms comparing the colocalization between DdMyo7-mCherry (x-axis) and GFP- DdMyo7 or GFP-DdMyo7 tail (y-axis) **E.** Analysis strategy for measuring entire cell peripheral intensity. **F.** Micrographs of cells expressing RFP-LifeAct, GFP-DdMyo7, GFP-Tail, or GFP- Motor-Forced Dimer (FD). **A-F** Scale bars are 10µm. **G.** Peripheral line scan intensity of cells from F. **H**. Sample cortical band intensity showing the mean and variation of intensities around the periphery. **I**. Cortical band standard deviation (SD; n>93 cells from 3 experiments for each group). A higher SD indicates asymmetric localization. One-way ANOVA with multiple comparison correct compared to actin, **** p<0.001, ns not significant.

### DdMyo7 is localized to dynamic cortical actin

The apparent asymmetrical distribution of DdMyo7 to the leading edge (Figure 1F) suggested that the actin at the cortex has a role in the recruitment and localization of DdMyo7. DdMyo7 and actin appear to be colocalized at the leding edge in cells expressing both GFP- DdMyo7 and an actin marker (Lifeact) (Figure 2A). Linescans all around the periphery show that indeed there is strong correlation of Lifeact and DdMyo7 fluorescence around the cell periphery (Figure 2B, yellow line from 2A). Cytofluorogram analysis also revealed strong correlation of the DdMyo7 and actin fluorescence intensities (Figure 2C). A line scan taken perpendicular to an extending leading edge showed a steady accumulation of DdMyo7 intensity at the same time as an increase in actin intensity (Figure 2D, E). DdMyo7 and actin normalized intensities during leading edge extension were measured for 10 independent cells and a spline fit shows a robust correlation with actin intensity (Figure 2F).

**Figure 2.**
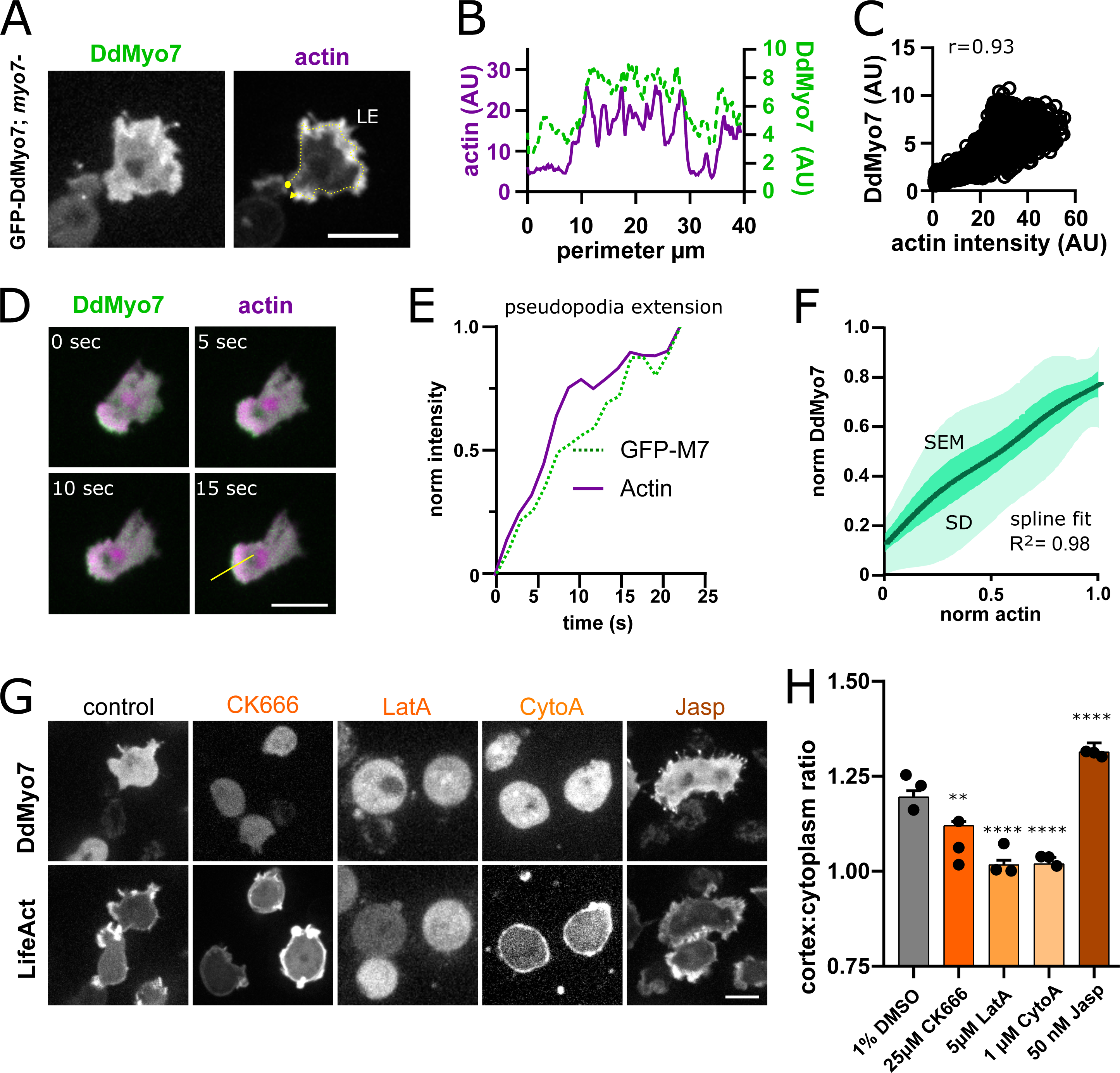
Actin dynamics regulate DdMyo7 recruitment to the cortex. **A**. *Dictyostelium* co-expressing GFP-DdMyo7 and RFP-Lifeact. **B**. Line intensity profile from yellow dotted line in A (circle=beginning, arrowhead indicates end of scan). **C**. Cytofluorogram showing the colocalization of actin and DdMyo7, r is correlation coefficient. **D.** Confocal image series of an extending pseudopod. **E.** Normalized linescan intensity profile of DdMyo7 and actin in extending pseudopod along line from panel D. **F.** Intensity correlation of GFP-DdMyo7 and RFP-LifeAct plotted as the average spline fit of 10 extending pseudopodia. **G.** Confocal micrographs of cells expressing GFP-DdMyo7 (top) or RFP-LifeAct (actin, bottom) under noted drug condition. **A,D,G.** Scale bar is 10µm. **H.** Cortex:cytoplasm ratio (cortex is 0.8µm band of cell periphery) of GFP-DdMyo7 of cells treated with anti-actin drugs, circles are experimental means. One-way ANOVA with multiple comparison correction, shown to 1% DMSO control, **p<0.01, p****<0.0001.

The dependence of DdMyo7 localization on actin polymerization at the cortex was tested by treating cells with actin modulating drugs. CytochalasinA (cytoA) binds to the fast-growing (barbed) end of actin filaments and blocks incorporation of actin monomers, capping and stabilizing filaments. LatrunculinA (latA) sequesters monomers and prevents actin filament growth, and CK-666 blocks the Arp2/3 complex and actin branching (Cooper, 1987; Coué et al., 1987; Nolen et al., 2009). Cells were incubated with each of these drugs and the impact on DdMyo7 cortical targeting was assessed by measuring the intensity ratio of DdMyo7 in a 0.8 µm band around the perimeter versus the cytoplasm (see Figure 1E). Control cells have a cortex:cytoplasm ratio of about ∼1.2, indicating an overall 20% enrichment of DdMyo7 on the cell cortex (Figure 2G, H; (Arthur et al., 2019)). Treatment of cells with either cytoA or latA reduced the ratio to ∼ 1 indicating a total loss of cortical localization of DdMyo7 (Figure 2H). CK-666 also significantly reduced DdMyo7 cortical recruitment (Figure 2G, H). Consistent with the reduction or loss of polymerized actin and cortical DdMyo7, filopodia formation was significant reduced with these actin modulating drugs (Table 1). Cells were also treated with Jasplakinolide (Jasp) that promotes monomer nucleation and stabilizes ADP-Pi actin filaments (Merino et al., 2018). Jasp treatment had the opposite effect, resulting in increased recruitment of DdMyo7 to the cortex and increased filopodia formation (Figure 2G, H; Table 1). The changes in DdMyo7 localization are specific to treatments that alter actin dynamics. The addition of either PI3Kinase inhibitors or microtubule depolymerizing agents such as nocodazole do not disrupt DdMyo7 cortical targeting (Supplemental Figure 2A, B). Control cells were imaged expressing GFP-CRAC (PIP3 marker, (Parent et al., 1998)), or GFP-tubulin (Neujahr et al., 1998), (Supplemental Figure 2A). The microtubule disrupting compound nocodazole had no effect on DdMyo7 in spite of the presence of two MF domains with micromolar affinity for microtubules in the DdMyo7 tail (Planelles-Herrero et al., 2016). Thus, the region of dynamic actin at the cortex controls the cortical recruitment of DdMyo7.

**Table 1.** Cortical recruitment ratio of DdMyo7 and filopodia per cell for GFP-DdMyo*7/myo7* null cells treated with various pharmacological compounds.

|  | Buffer Only | 1% DMSO |  |
| --- | --- | --- | --- |
| percent of cells with filopodia | 42 | 38 |  |
| filopodia number + SEM | 2.01±0.11 | 1.97±0.09 |  |
| cortex:cytoplasm ratio + SEM | 1.18±0.01 | 1.2±0.02 |  |
| N, n | 4, 238 | 3, 229 |  |
|  | 1µM<br>cytochalasinA | 5µM<br>cytochalasinA | 30µM<br>cytochalasinA |
| percent of cells with filopodia | 0 | 8.5 | 19 |
| filopodia number + SEM | 0 | 1.3±0.04 | 0.1 ±0.06 |
| cortex:cytoplasm ratio + SEM | 1.02±0.02 | 1.07±0.01 | 1.04±0.02 |
| N, n | 3, 66 | 3, 234 | 1, 42 |
|  | 1µM latrunculinA | 5µM latrunculinA | 15µM<br>latrunculinA |
| percent of cells with filopodia | 19 | 4.7 | 20 |
| filopodia number + SEM | 1.73±0.06 | 1.33±0.05 | 0.24±0.06 |
| cortex:cytoplasm ratio + SEM | 1.12±0.01 | 1.02±0.01 | 1.08 ± 0.02 |
| N, n | 4, 387 | 3, 127 | 1, 74 |
|  | 15nM<br>Jasplakinolide | 50nM<br>Jasplakinolide | 100nM<br>Jasplakinolide |
| percent of cells with filopodia | 55 | 72 | 48 |
| filopodia number + SEM | 1.47±0.23 | 3.89±0.2 | 2.21±0.09 |
| cortex:cytoplasm ratio + SEM | 1.19±0.03 | 1.34±0.03 | 1.24±0.02 |
| N, n | 1,84 | 3, 219 | 3, 337 |
|  | 20µM Ly294002 | 60µM Ly294002 | 2µM<br>Wortmannin |
| percent of cells with filopodia | 22 | 40 | 30 |
| filopodia number + SEM | 0.56±0.44 | 0.87±0.12 | 0.84±0.16 |
| cortex:cytoplasm ratio + SEM | 1.22±0.08 | 1.17±0.04 | 1.16±0.02 |
| N, n | 2,9 | 4,194 | 2,101 |
|  | 15µM Nocodazole | 50µM Nocodazole | 25µM CK666 |
| percent of cells with filopodia | n.c | 47.3 | 10.7 |
| filopodia number + SEM | n.c | 1.64±0.13 | 0.145±0.03 |
| cortex:cytoplasm ratio + SEM | 1.25±0.02 | 1.19±0.02 | 1.12±0.01 |
| N, n | 1,97 | 4, 169 | 3,366 |

### The role of VASP in DdMyo7 cortical recruitment

Filopodia initiation and extension is driven by regulators of actin polymerization such as VASP and formin ((Mattila and Lappalainen, 2008); Figure 3A,B). The actin bundler/polymerase DdVASP accumulates at the leading edge of cells and is important for filopodia formation (Han et al., 2002; Breitsprecher et al., 2008). Interestingly, the *Dictyostelium vasp* null mutant phenocopies the *myo7* null mutant - it lacks filopodia, has reduced adhesion and smaller cell size (Han et al., 2002; Tuxworth et al., 2001). This is of particular note as mammalian Myo10 and VASP are observed to co-transport to the tips of filopodia and co-immunoprecipitate, suggesting a role for Myo10 in the transport of VASP to filopodia tips to promote filopodia growth (Tokuo and Ikebe, 2004; Lin et al., 2013). There is currently no evidence that DdMyo7 and *Dictyostelium* VASP (DdVASP) interact with each other and co- immunoprecipitation experiments in *Dictyostelium* failed to detect any interaction (Supplemental Figure 3A). Surprisingly, in spite of this lack of interaction, DdMyo7 fails to target efficiently to the cortex of *vasp*- cells (Figure 3C, D). In contrast, DdVASP localizes to the cortex regardless of the presence of DdMyo7 (Figure 3C, D).

**Figure 3.**
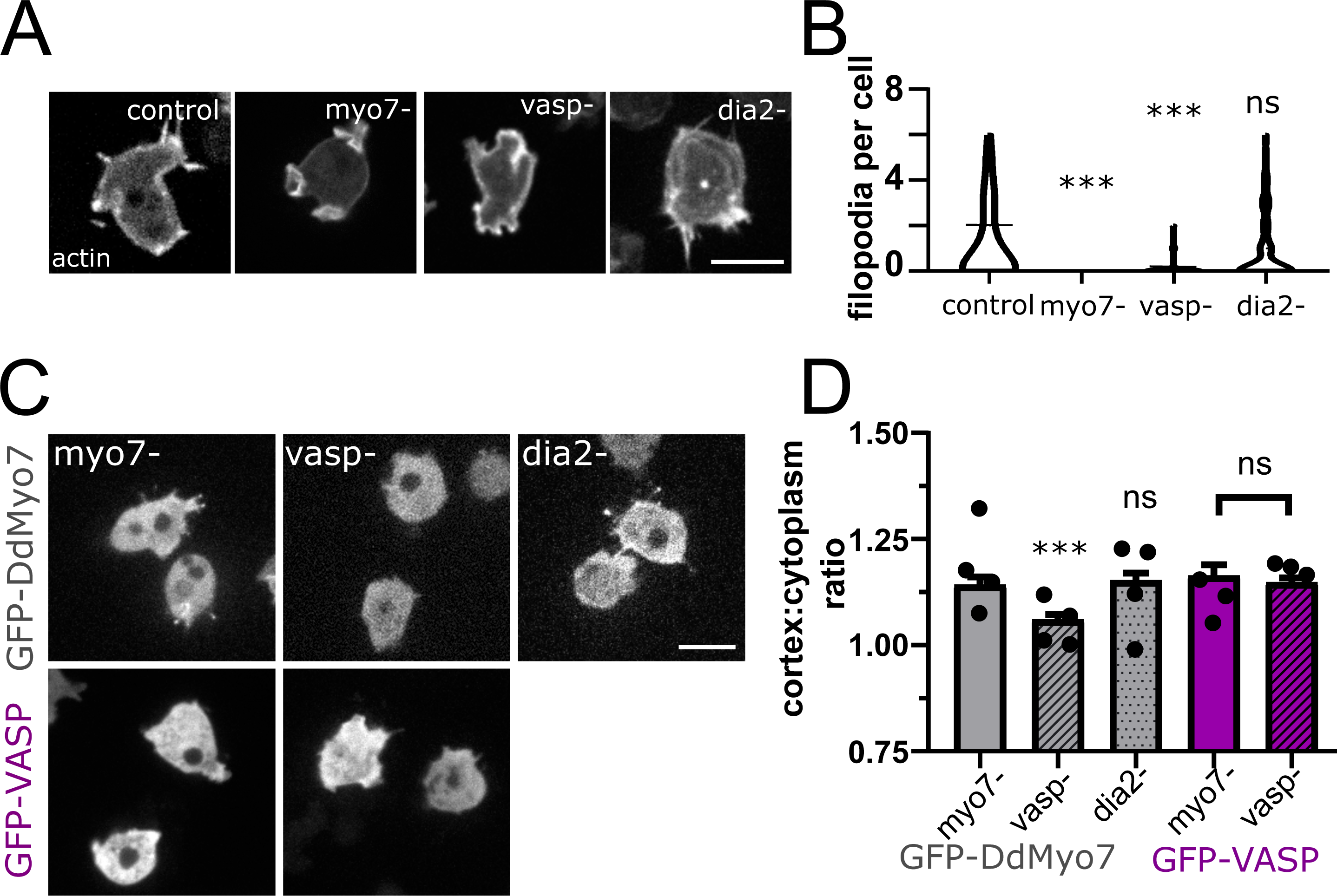
VASP is required for DdMyo7 cortical recruitment. **A**. Confocal images of wild type or *myo7* null, *vasp* null or *dia2* null cells expressing RFP-LifeAct (actin). **B.** Violin plot of number of filopodia per cell. **C.** Micrographs of cells expressing GFP-DdMyo7 (top) or GFP-VASP in *myo7* null*, vasp* null or *dDia2* null cells. **D**. Quantification of the cortical band (0.8µm of periphery) relative to the cytoplasmic intensity of either GFP-Myo7 or GFP-VASP. **A,C** Scale bar is 10µm. **B, D,** One-way ANOVA with multiple comparison correction, ns, not significant, p***<0.001, circles are experimental means.

Formins are localized to the leading edge of migrating cells and play an important role in filopodia formation (Schirenbeck et al., 2005; Pellegrin and Mellor, 2005; Yang et al., 2007). Formins promote actin polymerization by incorporating actin monomers at the barbed end of actin filaments and elongate parallel actin filaments (Breitsprecher and Goode, 2013; Mellor, 2010). The *Dictyostelium* diaphanous related formin dDia2 makes significant contributions to filopodia formation and is required for normal filopodia length (Figure 3A, B; (Schirenbeck et al., 2005)). Formin activity is not required for cortical recruitment of DdMyo7 as it is found to localize normally to the cortex of *dDia2* null (Figure 3C, D). These data reveal that DdVASP or its actin polymerization activity is critical for DdMyo7 recruitment and suggests that this bundler/polymerase acts upstream of DdMyo7.

### Selective recruitment of DdMyo7 requires specific actin polymerization factors

The finding that DdVASP is critical for DdMyo7 cortical targeting raises the question of how it is acting to recruit this myosin. One simple explanation is that DdMyo7 is targeting to regions of the cell cortex where dynamic actin polymerization occurs. This hypothesis was tested by examining if robust recruitment of DdMyo7 could occur in the absence of DdVASP when actin polymerization was stimulated using different manipulations. Treatment of *Dictyostelium* with LatA treatment followed by washout produces robust actin waves (Gerisch et al., 2004). These waves are made of a dense Arp2/3 branched actin meshwork and class I myosins (MyoB, MyoC) (Jasnin et al., 2019; Brzeska et al., 2020). Cells expressing both RFP-LimE (a marker for F-actin) and GFP-DdMyo7 were induced to form waves that are readily apparent as broad circles of actin that emanate outwards towards the cell periphery. Actin waves were generated in either control or *vasp* null cells and in all cells visualized (n=6 per genotype) all waves are completely devoid of DdMyo7 (Figure 4A). Next, *vasp-* cells were treated with Jasp to stimulate actin polymerization (Supplemental Figure 4A). No increase in cortical targeting of DdMyo7 was observed in *vasp* nulls treated with Jasp when compared to untreated cells (Figure 4B, E). Blocking capping protein by overexpressing the capping protein inhibitor V-1 stimulates filopodia formation in *Dictyostelium* (Supplemental Figure 4B, (Jung et al., 2016)). V-1 also stimulates cortical actin network formation in *vasp* null *Dictyostelium* (Figure 4C). No increase in targeting was observed in *vasp* nulls with induced overexpression of V-1 (Figure 4B, D), nor was filopodia formation restored (Figure 4F).

**Figure 4.**
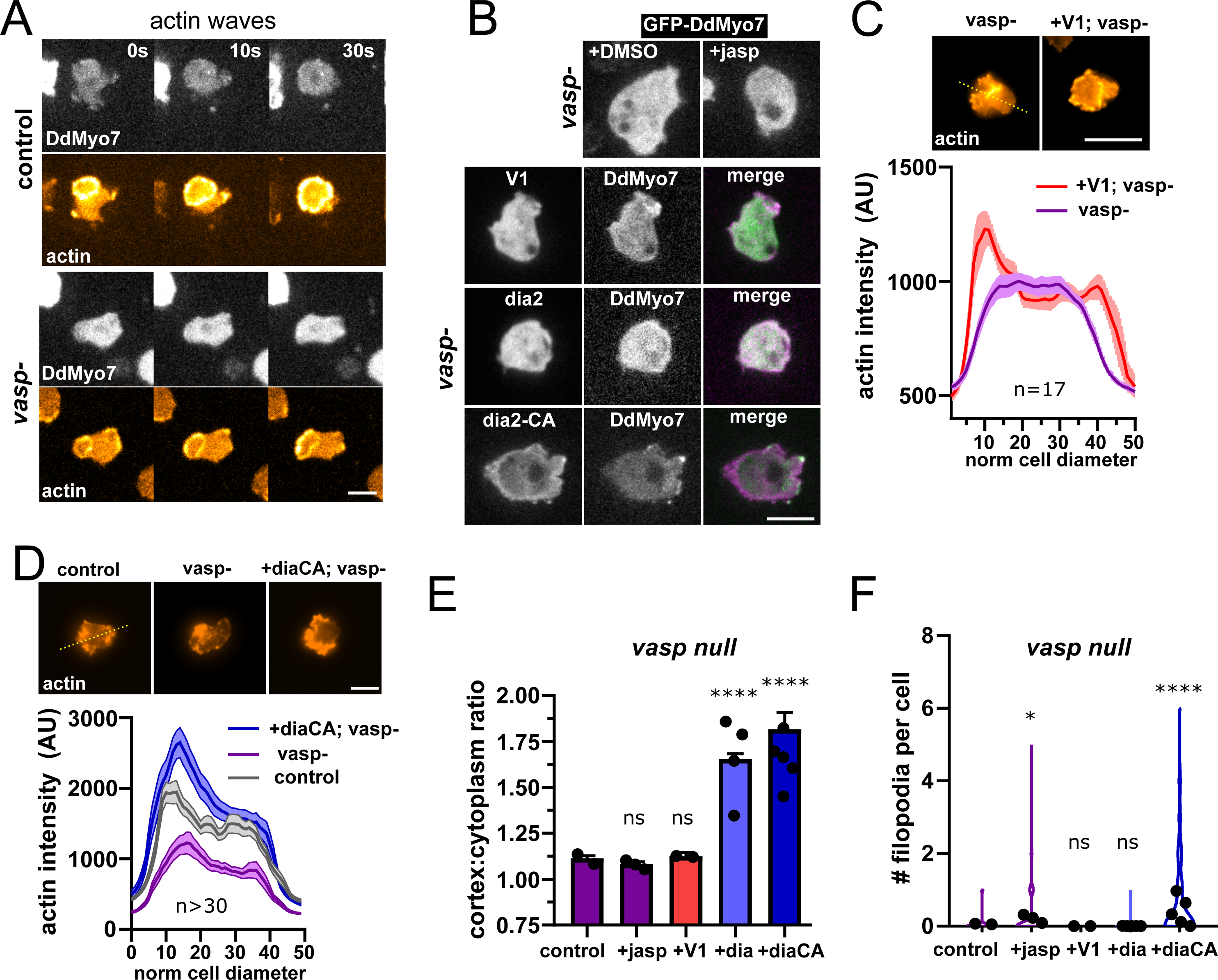
Linear actin polymerization drives DdMyo7 to the cortex. **A.** Images series showing DdMyo7 is absent from LatrunculinA-induced actin waves in control (top) or *vasp null* (bottom) cells. **B.** (top) Confocal images of GFP-DdMyo7 in *vasp* null cells treated with either DMSO or 50nM Jasp treatment. (bottom) Images of cells expressing DdMyo7 and different actin modulating proteins. **C, D.** Average actin intensity (phalloidin staining, top) of cells through the longest cell axis. The line is the mean and the shaded area is the SEM (graphs, bottom). **A-D.**, Scale bar is 10µm. **E.** Quantification of the cortical band intensity of DdMyo7 in *vasp* null cells, with no treatment, treated with Jasp, or also overexpressing V-1, dia2, or dia2-CA. **F.** Violin plot of filopodia per cell**. E-F.** One-way ANOVA with multiple comparison correction, ns, not significant, * p<0.05, * p****<0.0001.

Activation of formins can also result in robust actin polymerization. These polymerases are tightly controlled by autoinhibition with the conserved DAD domain binding to the N-terminal DID domain, blocking actin nucleation activity. Mutation of conserved basic residues in the C-terminal DAD domain creates a constitutively activated (CA) formin (Wallar et al., 2006). A pair of charged residues are present in the dDia2 DAD domain and these were mutated - R1035A, R1036A - to create a CA formin (see alignment in Supplemental Figure 4C). Either dDia2 or dDia2-CA overexpressing *vasp* null cells exhibit restored and increased DdMyo7 cortical targeting (Figure 4B, E). Overexpression of dDia2-CA also promoted a modest rescue of filopodia formation (Figure 4F; Table 2). Linescans of cells stained with phalloidin show that while *vasp* nulls have less cortical F-actin than wild type cells, expression of dDia2- CA in *vasp* nulls resulted in increased cortical F-actin (Figure 4D). In summary, stimulation of actin polymerization by inducing actin waves, treating cells with Jasp, or blocking capping protein is not sufficient to promote DdMyo7 cortical recruitment in *vasp* null cells. In contrast, expression of activated formin is sufficient for cortical recruitment in *vasp* null cells. This strongly suggests that DdMyo7 cortical targeting specifically requires the activity of actin regulators that generate parallel actin filaments.

**Table 2.**
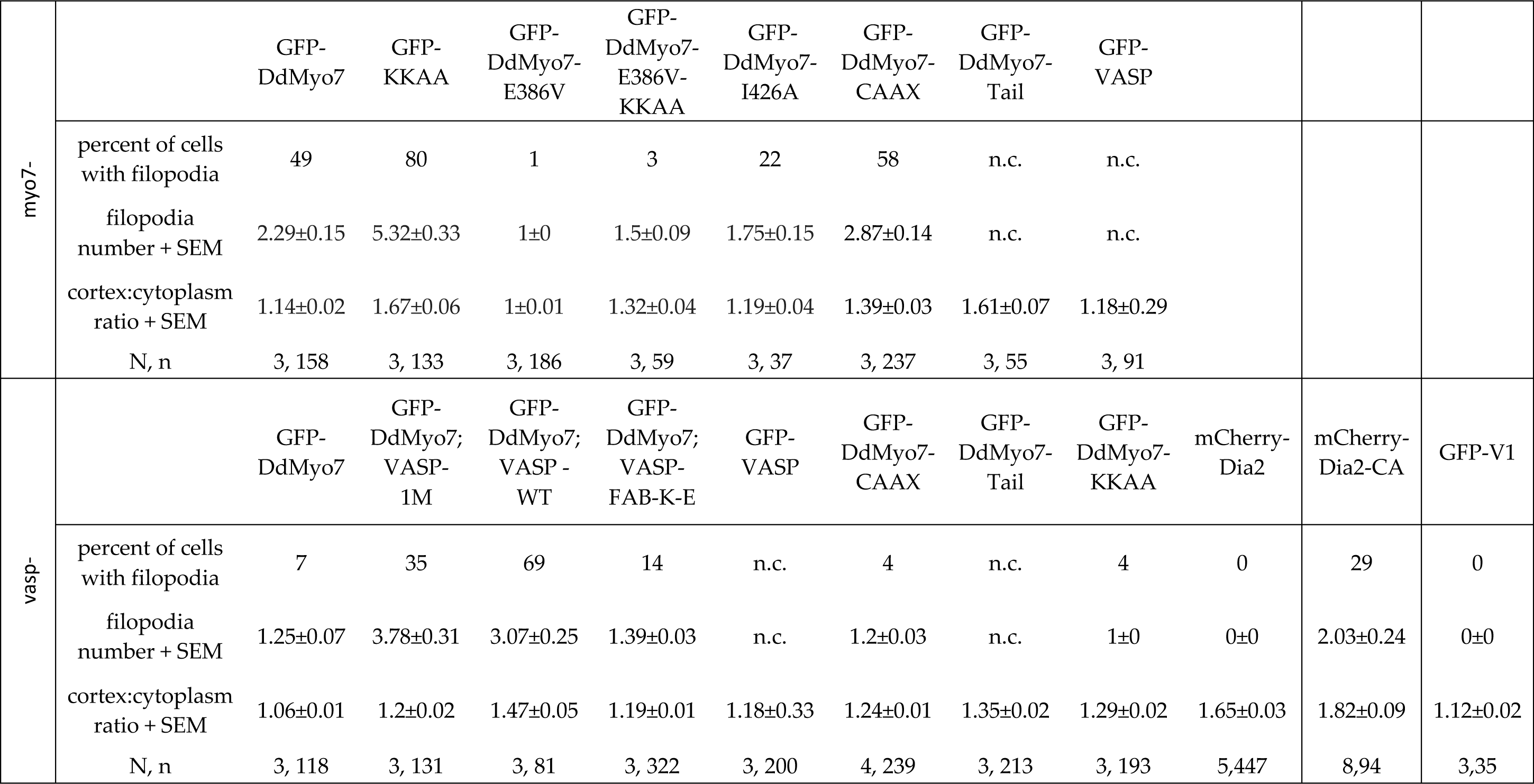
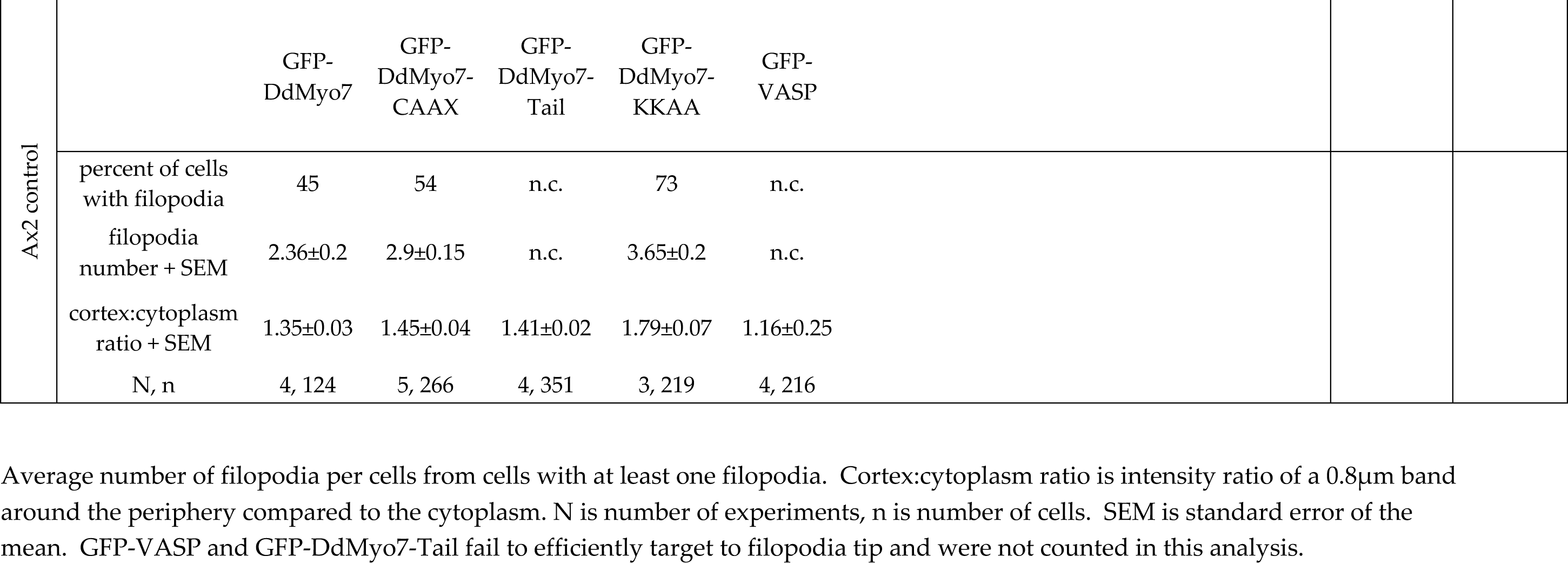
Quantification of filopodia number and cortical targeting.

| myo7- |  | GFP-DdMyo7 | GFP-KKAA | GFP-DdMyo7-E386V | GFP-DdMyo7-E386V-KKAA | GFP-DdMyo7-I426A | GFP-DdMyo7-CAAX | GFP-DdMyo7-Tail | GFP-VASP |  |  |  |
| --- | --- | --- | --- | --- | --- | --- | --- | --- | --- | --- | --- | --- |
|  | percent of cells<br>with filopodia | 49 | 80 | 1 | 3 | 22 | 58 | n.c. | n.c. |  |  |  |
|  | filopodia<br>number + SEM | 2.29±0.15 | 5.32±0.33 | 1±0 | 1.5±0.09 | 1.75±0.15 | 2.87±0.14 | n.c. | n.c. |  |  |  |
|  | cortex:cytoplasm<br>ratio + SEM | 1.14±0.02 | 1.67±0.06 | 1±0.01 | 1.32±0.04 | 1.19±0.04 | 1.39±0.03 | 1.61±0.07 | 1.18±0.29 |  |  |  |
|  | N, n | 3, 158 | 3, 133 | 3, 186 | 3, 59 | 3, 37 | 3, 237 | 3, 55 | 3, 91 |  |  |  |
| vasp- |  | GFP-DdMyo7 | GFP-DdMyo7;<br>VASP-1M | GFP-DdMyo7;<br>VASP -<br>WT | GFP-DdMyo7;<br>VASP-FAB-K-E | GFP-VASP | GFP-DdMyo7-CAAX | GFP-DdMyo7-Tail | GFP-DdMyo7-KKAA | mCherry-Dia2 | mCherry-Dia2-CA | GFP-V1 |
|  | percent of cells<br>with filopodia | 7 | 35 | 69 | 14 | n.c. | 4 | n.c. | 4 | 0 | 29 | 0 |
|  | filopodia<br>number + SEM | 1.25±0.07 | 3.78±0.31 | 3.07±0.25 | 1.39±0.03 | n.c. | 1.2±0.03 | n.c. | 1±0 | 0±0 | 2.03±0.24 | 0±0 |
|  | cortex:cytoplasm<br>ratio + SEM | 1.06±0.01 | 1.2±0.02 | 1.47±0.05 | 1.19±0.01 | 1.18±0.33 | 1.24±0.01 | 1.35±0.02 | 1.29±0.02 | 1.65±0.03 | 1.82±0.09 | 1.12±0.02 |
|  | N, n | 3, 118 | 3, 131 | 3, 81 | 3, 322 | 3, 200 | 4, 239 | 3, 213 | 3, 193 | 5,447 | 8,94 | 3,35 |

| Ax2 control |  | GFP-DdMyo7 | GFP-DdMyo7-CAAX | GFP-DdMyo7-Tail | GFP-DdMyo7-KKAA | GFP-VASP |
| --- | --- | --- | --- | --- | --- | --- |
|  | percent of cells with filopodia | 45 | 54 | n.c. | 73 | n.c. |
|  | filopodia number + SEM | 2.36±0.2 | 2.9±0.15 | n.c. | 3.65±0.2 | n.c. |
|  | cortex:cytoplasm ratio + SEM | 1.35±0.03 | 1.45±0.04 | 1.41±0.02 | 1.79±0.07 | 1.16±0.25 |
|  | N, n | 4, 124 | 5, 266 | 4, 351 | 3, 219 | 4, 216 |

### Mechanism of VASP mediated actin polymerization required for DdMyo7 recruitment

The properties of VASP critical for DdMyo7 recruitment were investigated using separation of function mutations. VASP accelerates the rate of actin polymerization, bundles actin filaments and blocks capping protein from binding to the plus ends of actin filaments (Breitsprecher et al., 2008, 2011; Hansen and Mullins, 2010). VASP is a constitutive tetramer that can bundle actin by binding to the sides of filaments (see Figure 5A; (Bachmann et al., 1999; Breitsprecher et al., 2008; Brühmann et al., 2017). DdVASP F-actin binding is mediated by a region within the EVH2 domain (aa 264-289) and tetramerization by a region at the C-terminus (aa 341-375) (Schirenbeck et al., 2006; Breitsprecher et al., 2008). Two different mutants were used to test if the VASP bundling activity is required for DdMyo7 recruitment (Figure 5D). A monomeric VASP was created by deleting the tetramerization domain (1M; Δ341-375)) and an F-actin binding mutant was generated in which conserved lysines were mutated to glutamate (FAB K-E; KR275,276EE + K278E + K280E; based on (Hansen and Mullins, 2010)). The FAB K-E mutations are predicted to slow but not eliminate actin polymerization (Breitsprecher et al., 2008; Schirenbeck et al., 2006; Applewhite et al., 2007; Hansen and Mullins, 2010). Co-expression of either DdVASP-1M or DdVASP-FAB K-E with DdMyo7 in *vasp* null cells cannot fully restore DdMyo7 cortical recruitment (Figure 5B, E). The monomeric VASP-1M mutant partially rescues filopodia formation while the FAB K-E mutant does not (Figure 5C). The residual filopodia forming activity of the VASP-1M monomer could be attributed to its remaining anti-capping activity that is lost in the F-actin binding mutant (Breitsprecher et al., 2008; Hansen and Mullins, 2010). Both mutants are predicted to lack any bundling activity and show decreased recruitment of DdMyo7 to the cortex. These data indicate that bundling linear F-actin at the membrane could promote DdMyo7 targeting.

**Figure 5.**
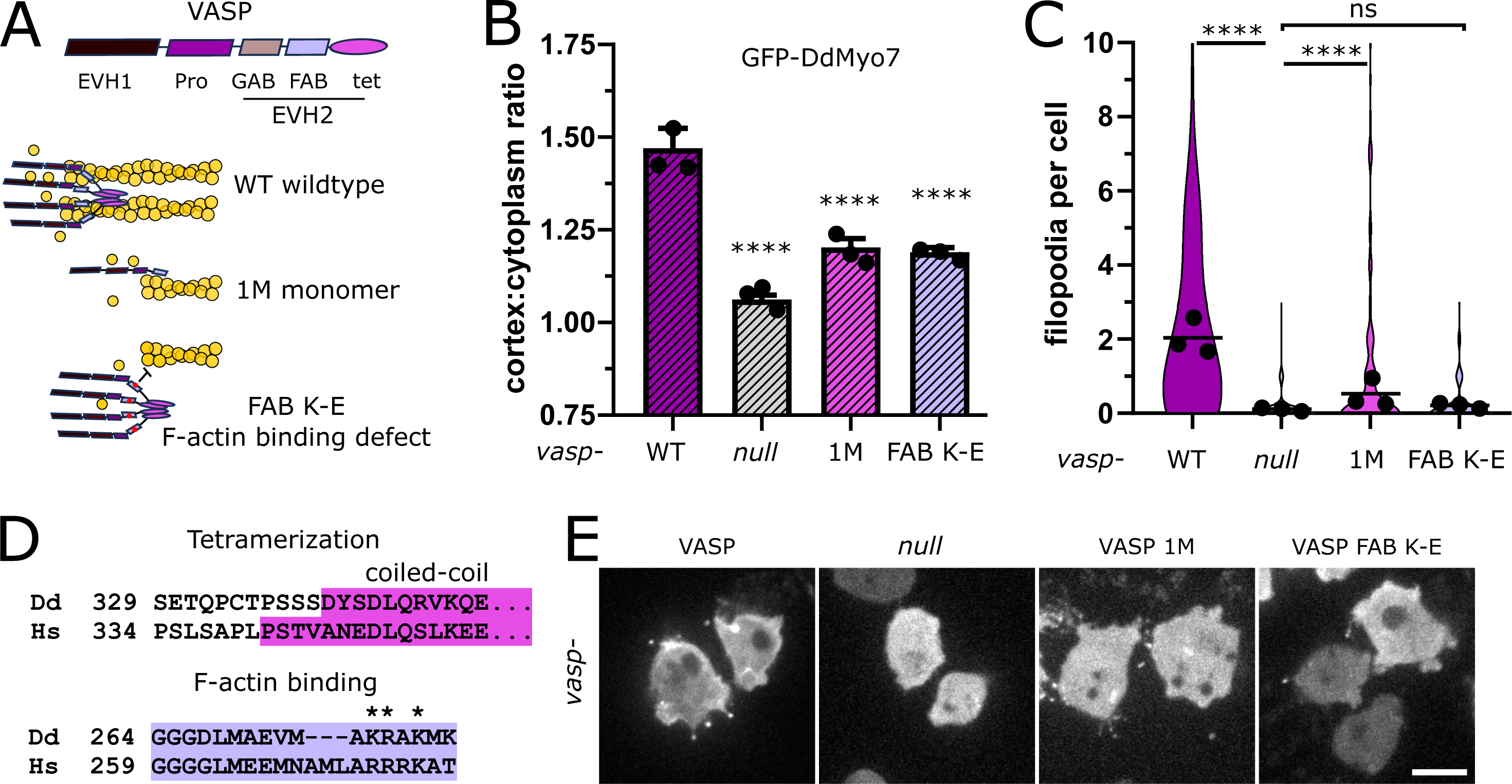
Reduced cortical recruitment of DdMyo7 by VASP mutants. **A**. Schematic of domains of DdVASP (top) and proposed interaction of DdVASP wildtype, monomeric, and F-actin binding (FAB K-E) mutant with actin filaments. **B.** Quantification of the cortical recruitment of DdMyo7 co-expressed in the *vasp* null with wildtype or mutant DdVASP rescue constructs. **C**. Quantification of filopodia per cell of *vasp* null cells with wildtype or mutant DdVASP rescue constructs. **B-C**. Circles represent experimental means. One-way ANOVA with multiple comparison correction, p****<0.0001, ns not significant. **D.** Clustal Omega alignment of *Dictyostelium* and human VASP with conserved domains highlighted and mutated residues starred. **E.** Micrographs of GFP-DdMyo7 in *vasp* nulls, or *vasp* nulls expressing wildtype DdVASP or mutant DdVASP rescue constructs. Scale bar is 10µm.

### Release of DdMyo7 autoinhibition dictates the spatial restriction of its recruitment

DdMyo7 is enriched at the leading edge of cells, in contrast to the tail alone that localizes all around the cortex (Figure 1F, I). This difference suggests that head-tail autoinhibition restricts DdMyo7 localization. DdVASP is required for cortical localization but the lack of interaction between DdMyo7 and DdVASP suggests an indirect role. Furthermore, the GFP-DdMyo7 tail domain in *vasp* null cells targets to the cortex efficiently (Figure 6A-C). This confirms that direct DdMyo7 tail - DdVASP binding is not required for localization. Together these results suggested that VASP mediated actin networks release autoinhibition of DdMyo7. If so, then loss of autoinhibition regulation of DdMyo7 should eliminate the requirement of VASP for DdMyo7 to target to the cortex. To test this, two highly conserved charged residues at the extreme C-terminus of DdMyo7 (K2333, K2336) essential for head-tail autoinhibition were mutated (Yang et al., 2009; Petersen et al., 2016). Overexpression of this constitutively active mutant (DdMyo7- KKAA) stimulates filopodia formation and increases cortical localization in wild type cells (Table 2, (Petersen et al., 2016; Arthur et al., 2019)). DdMyo7-KKAA was expressed in *vasp* null cells where it targets to the cortex (Figure 6A-B). The cortical targeting of DdMyo7-KKAA in *vasp* nulls is not as robust as seen in control cells (Figure 6C), and it is localized more uniformly around the cortex in *vasp* nulls (less enriched in the leading edge or pseudopod) compared to controls (Figure 6E). These observations are consistent VASP activity contributing to shift the equilibrium of the autoinhibited myosin to the open, active state, recruiting the activated DdMyo7 to the cortex.

**Figure 6.**
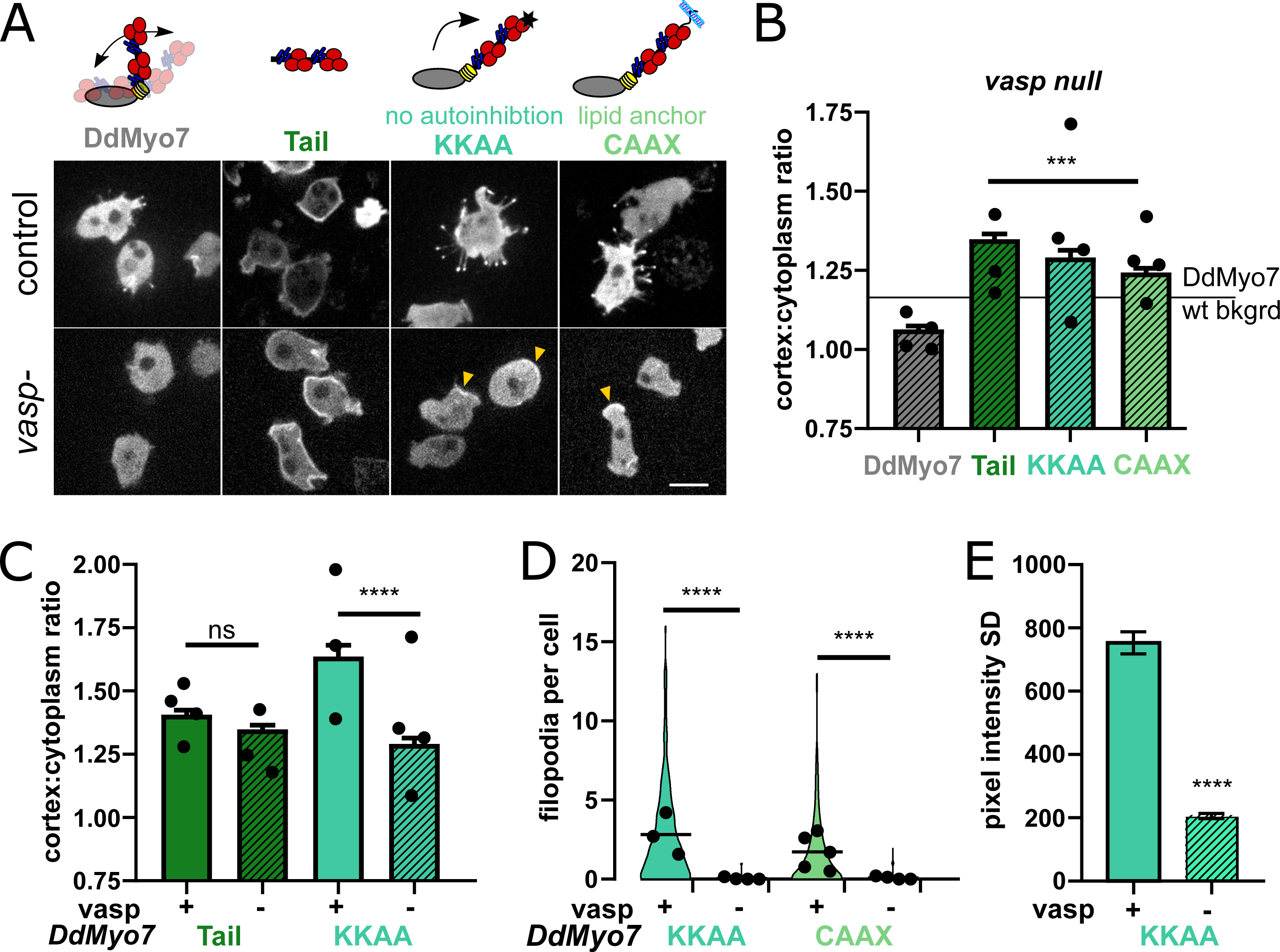
VASP relieves DdMyo7 head-tail autoinhibition to promote targeting and filopodia formation. **A**. (top) Diagrams depicting mutants analyzed. (bottom) Micrographs of GFP- DdMyo7 fusion proteins in control and *vasp* null cells, scale bar is 10µm **B.** Quantification of cortical recruitment of GFP-DdMyo7 and variants in *vasp^-^* cells. The line represents the mean GFP-DdMyo7 recruitment in wild type cells. **C.** Comparison of cortical targeting of activated KKAA or tail in *vasp* null versus control cells. **D.** Quantification of number of filopodia per cell in control or *vasp* null cells. **B-D.** Circles represent experimental means. One way ANOVA with multiple comparison test, ns not significant, p***<0.001, p****<0.0001, ns, not significant. **E.** Quantification of the cortical band intensity variation of DdMyo7-KKAA in control versus vasp null cells. Students t-test ****p<0.0001.

### VASP and cortical DdMyo7 are both required for filopodia formation

The finding that loss of head-tail autoinhibition restores cortical recruitment of DdMyo7 raised the question of whether an autoactivated motor is sufficient to rescue the filopodia formation defect seen in the *vasp* null cells. Expression of DdMyo7-KKAA in *vasp* nulls did not rescue their filopodia formation defect (Figure 6D), indicating that a specific DdVASP activity is required for filopodia initiation. The requirement for both cortical DdVASP and DdMyo7 for filopodia formation was further tested by targeting of the myosin to the cortex or membrane by fusing a prenylation sequence to its C-terminus (DdMyo7-CAAX, adapted from (Weeks et al., 1987)). DdMyo7-CAAX was robustly localized to the cortex in *myo7* null, wild type and *vasp* null cells (Fig. 6A, B; Table 2). Expression of DdMyo7-CAAX in wildtype or *myo7* nulls cells significantly stimulated filopodia formation (Figure 6A, D, Table 2). However, filopodia are not formed when DdMyo7-CAAX was expressed in *vasp* null cells (Figure 6A, D). These results establish that targeting DdMyo7 to the membrane alone is not sufficient for filopodia initiation and that DdVASP must also be present.

### Role of myosin motor activity in targeting and filopodia formation

The data suggest that activation of autoinhibited DdMyo7 is promoted at the front of the cell in regions rich in newly polymerized F-actin. Actin binding by myosin involves conformational changes in the myosin head (Houdusse and Sweeney, 2016) and could destabilize the interactions between the motor head and the tail. To test this, two DdMyo7 mutants were designed, each with mutations in highly conserved connectors of the motor domain. Their functions are established by studies performed on the highly conserved motor domain DdMyo2 (Sasaki et al., 2003; Friedman et al., 1998). The non-hydrolyzer mutant (E386V) binds ATP but cannot hydrolyze it, and thus stays in weak actin binding state, while the uncoupler mutant (I426A) can undergo conformational changes allowing strong interactions with F-actin but these conformational changes are not transmitted to the lever arm to produce force (illustrated in Figure 7A).

**Figure 7.**
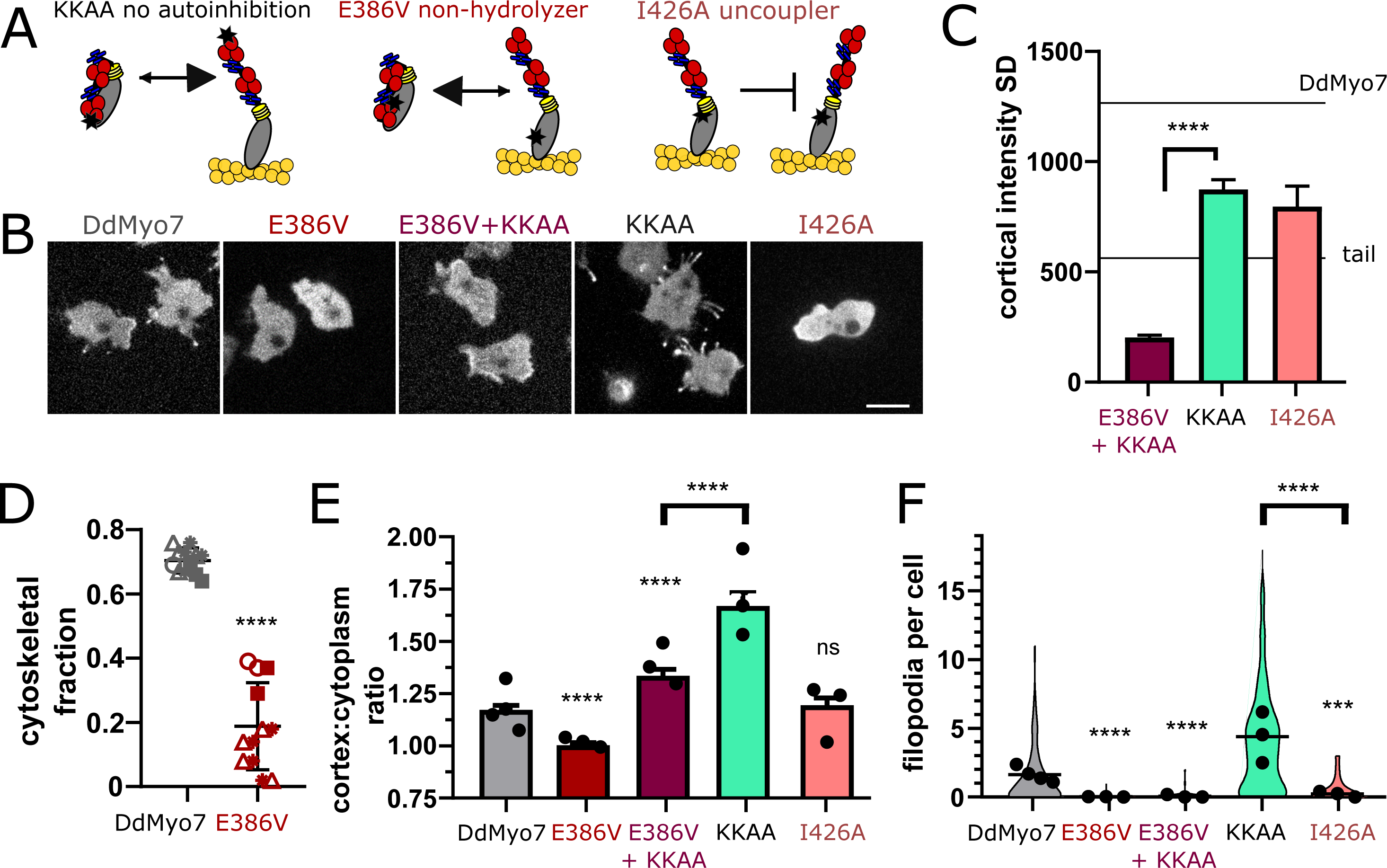
DdMyo7 motor activity is required to release autoinhibition. **A.** Schematic of proposed effect of mutations on DdMyo7 function. **B.** Confocal images of *myo7* null cells expressing GFP-DdMyo7 fusion proteins, scale bar is 10µm. **C.** Quantification of the cortical band intensity variation. Mean lines from Figure 1I data on graph for comparison. DdMyo7 versus I426A uncoupler, p**<0.01. **D.** Fraction of DdMyo7 cosedimenting with the cytoskeleton, symbols with the same shape are technical replicates, students t-test ****p<0.0001. **E.** Quantification of cortical recruitment of DdMyo7 and mutants. **F.** Filopodia number per cell of wildtype and mutant DdMyo7. **E-F.** Data for KKAA is taken from Figure 6, experimental means shown as circles. **C, E, and F**, One-way ANOVA with multiple comparison correction, p***<0.001, p****<0.0001, ns not significant.

The non-hydrolyzer (E386V) has a mutated glutamate in Switch II that is required to assist hydrolysis (Supplemental Figure 5A, (Friedman et al., 1998)). The non-hydrolyzer failed to efficiently cosediment with the actin cytoskeleton upon centrifugation (Figure 7D, Supplemental Figure 5B) indicating that it was indeed a weak actin binding mutant. The DdMyo7-E386V mutant does not target efficiently to the cell periphery (Figure 7B, E) or rescue filopodia formation (Figure 7F; Table 2). The lack of cortical localization by the non-hydrolyzer suggests that the tail remains bound to the motor, favoring an autoinhibited conformation, and is not free to bind to the cortex (see Figure 7A). To test this, the autoinhibition point mutations (KKAA) were introduced into the non-hydrolyzer to disrupt the head/tail interface. Blocking autoinhibition (KKAA) in the non-hydrolysis mutant (DdMyo7-E386A + KKAA) strikingly restores cortical targeting (Figure 7B, E; Table 2) but not filopodia formation (Figure 7F; Table 2).

A second motor mutation, I426A, resides in the relay helix and disrupts the interface with the converter that is critical to direct the swing of the lever arm during force generation. This mutant uncouples ATP hydrolysis from force generation (Supplemental Figure 5A, (Sasaki et al., 2003)). Thus, the uncoupler (DdMyo7-I426A) undergoes actin-activated ATP hydrolysis, likely due to conformational changes directed from binding to actin, but it cannot exert force (illustrated in Figure 7A). DdMyo7-I426A cortical targeting is similar to what is seen for wild type DdMyo7 (Figure 7B, E). This indicates that actin binding destabilizes the autoinhibited form in this mutant. DdMyo7-I426A fails to efficiently rescue the filopodia formation defect of *myo7* null cells despite its targeting to the cortex (Figure 7F and Table 2), establishing that force generation by DdMyo7 is essential for filopodia initiation.

The uncoupler mutant motor is predicted to have normal actin binding and indeed its cortical asymmetry is similar to wildtype DdMyo7 (Figure 7B, C). In contrast, the motor of the non-hydrolyzer DdMyo7-E386A + KKAA mutant has weak actin binding so interaction with the cortex would be directed by the tail. As predicted, the DdMyo7-E386A + KKAA mutant is localized uniformly around the cortex (low cortical SD, Figure 7B, C). These observations provide support for the model that while the tail aids overall cortical localization, the motor- actin binding refines leading edge targeting of DdMyo7.

## Discussion

Filopodia formation in amoeba requires both a MF myosin and VASP (Tuxworth et al., 2001; Han et al., 2002). The motor domain of the MF myosin DdMyo7, not the tail, targets it to actin-rich dynamic regions of the cortex where filopodia emerge (Figure 1). Actin network dynamics generated by DdVASP activity *in vivo* are important for myosin localization, promoting release of head-tail autoinhibition (Figure 6), freeing and activating the motor domain for binding to actin at the leading edge (Figures 1 and 7). This controlled activation of MF myosin leads to tuned activity at the right time and the right place to concentrate motors and facilitate filopodia formation, mainly from the leading edge of the cell. The role of DdVASP goes beyond simple recruitment of the motor since targeting of DdMyo7 at the membrane is not sufficient for filopodia initiation in *vasp* null cells (Figure 6A, D). Thus, both MF myosin and DdVASP together are needed to reorganize and polymerize actin at the cell cortex for filopodia to form (Figure 3 and 4). The data presented here support a model that first VASP activity, possibly bundling or polymerizing unbranched actin at the leading edge is upstream of DdMyo7 activation (Figure 8A,B). DdMyo7 is activated by motor binding to a DdVASP generated actin network (Figure 8C), specifically the open state (Figure 8E) is stabilized by motor binding to the linear, bundles of actin resulting from DdVASP activity. Together, DdVASP and DdMyo7 drive filopodia initiation by further bundling and polymerizing actin (Figure 8D).

**Figure 8.**
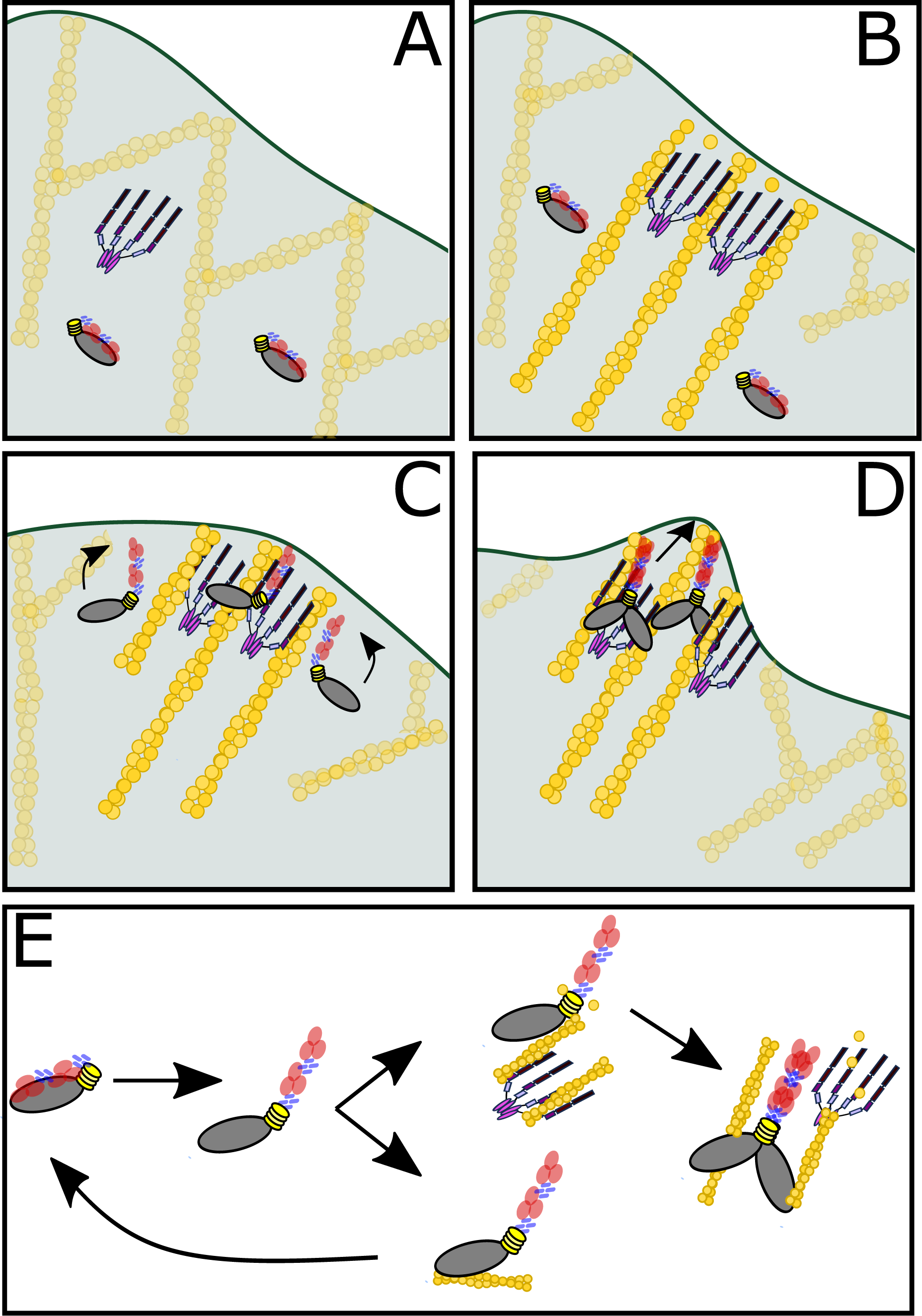
Model of DdMyo7 and VASP mediated filopodia initiation. **A.** The leading edge of the cell has a branched actin network. **B**. VASP polymerizes actin at the leading edge, organizing the filaments into linear, parallel bundles. **C**. Autoinhibited DdMyo7 is activated and the motor domain binds to actin within the VASP-actin network. **D**. Cooperative actions of VASP (bundles and polymerizes) and DdMyo7 dimers (bundle) organized actin filaments into nascent filopodia that continue to elongate by actin polymerization. **E**. Proposed model of DdMyo7 motor activation by actin. (left) depiction of the conformation change when the myosin is activated by relief of head-tail autoinhibition. (right) depiction of activated DdMyo7 binding actin via its motor domain, followed by dimerization and bundling of actin, generating force to bring actin filaments together. If the motor encounters a VASP actin network, it can dimerize and generate force, but if not, then it cycles back to an off state.

### The role of DdVASP in myosin recruitment

Disruption of the dynamic actin network at the cortex that occurs by loss of *vasp* (Figure 3B, Figure 4D) or by treatment with anti-actin drugs such as cytoA or latA (Figure 2G, H) results in loss of cortical localization of DdMyo7. *Dictyostelium* VASP is a potent actin polymerase, its activity speeds actin elongation, bundles filaments and blocks capping proteins from binding to the growing ends of actin filamments (Breitsprecher et al., 2008). Both its actin polymerization and bundling activities could be important for DdMyo7 recruitment (Figure 5). In the absence of DdVASP, increased cortical actin generated by the formin dDia2 can also recruit and activate DdMyo7 (Figure 4B - E). However, recruitment is not restored by other means of increasing actin polymerization, including generating actin waves, polymerizing actin with Jasp or blocking capping protein by overexpression of V-1. Thus, DdVASP does not simply recruit DdMyo7 by generating new actin polymers, but rather it is the nature of the network that DdVASP builds, likely growing linear actin filaments that are bundled together in parallel, that is critical for activation of DdMyo7.

The actin cortex of *vasp* nulls is less dense than found for wild type cells as revealed by phalloidin staining, suggesting that a robust leading edge actin network collapses without VASP (Figure 4D). Interestingly, evidence for VASP-dependent changes in the cortical actin network has recently been shown in B16 melanoma cells (Damiano-Guercio et al., 2020). The cortical actin network is normally dense and with many filaments oriented perpendicular to the membrane but in the absence of VASP, the network is more disperse and actin filaments are oriented in shallower angles with respect to the membrane (Damiano-Guercio et al., 2020). Thus, the formation of a VASP-mediated actin network at the cortex is likely to be important in this context as well as in other cell types.

### Actin-dependent release of autoinhibition and leading edge targeting

The motor domain of DdMyo7 is responsible for localizing this myosin to the dynamic leading edge of the cell, not the tail (Figures 1 and 2). The motor is typically sequestered by head-tail autoinhibition, a widely used mechanism for controlling myosin activity and reducing unnecessary energy expenditure (Heissler and Sellers, 2016). Overcoming autoinhibition is thus essential for liberating the motor and allowing it to target DdMyo7 to the dynamic cortex. DdMyo7 head/tail autoinhibition is mediated by charged residues in the FERM domain, similar to fly myosin 7a, and this regulates both cortical targeting and filopodia number (Petersen et al., 2016; Arthur et al., 2019; Yang et al., 2009). Blocking autoinhibition results in enhanced cortical targeting and filopodia formation, and bypasses the loss of cortical targeting observed in the absence of DdVASP (Figure 6). Motor activity (i.e. ATP hydrolysis) and actin binding are needed to release autoinhibition and promote cortical recruitment of DdMyo7 (Figure 7). Weak actin binding, even in the absence of autoinhibition does not promote leading edge targeting. In contrast, a mutant that can bind actin but uncouples force generation from ATP hydrolysis is localized similarly to wild type DdMyo7 (Figure 7A-C). The uniform localization of the tail all around the cortex suggests that while the tail provides general cortical targeting, an active motor refines where activation occurs via its specific recognition of a dynamic actin-rich leading edge of the cell that is generated by DdVASP activity. Thus, motor domain mediated binding to a specific actin network is a key way to restrict localization of DdMyo7 during filopodia formation.

### Conserved and divergent models of filopodia myosin function

The mechanism of autoinhibition via motor-tail stabilizing interactions has been conserved throughout the evolution of myosins (Umeki et al., 2011; Sakai et al., 2011; Yang et al., 2009; Petersen et al., 2016; Weck et al., 2017; Heissler and Sellers, 2016). Mechanisms to overcome autoinhibition likely developed to restrict the recruitment and activation of myosins in time and space, allowing cells to fine-tune their activity. Here it is shown that DdMyo7 autoinhibition is relieved where the myosin will act by the cortical actin network that is generated by DdVASP. While this differs from the PIP3-mediated recruitment of metazoan Myo10, once activated these two evolutionarily distant filopodia myosins use similar mechanisms to generate filopodia. They both dimerize upon recruitment to the actin-rich cortex (Arthur et al., 2019; Lu et al., 2012) where they likely contribute to the reorganization of the Arp2/3 branched actin network to orient actin filaments perpendicular to the membrane (Svitkina et al., 2003; Tuxworth et al., 2001; Berg and Cheney, 2002; Tokuo et al., 2007; Arthur et al., 2019). Interestingly, both DdMyo7 and Myo10 work in cooperation with VASP, although they appear to do so in different ways. In the case of Myo10, it is seen to co-transport with VASP along the length of filopodia during extension and co-immunoprecipitate with VASP, although it is not known if the two proteins interact directly in vivo (Tokuo and Ikebe, 2004; Kerber et al., 2009; Lin et al., 2013). In contrast, the activity of DdVASP generates an actin-rich leading edge that recruits and activates DdMyo7 and there is no evidence that the two proteins interact with each other at present (Figures 1, 3 and 6; Supplementary Figure 3).

*Dictyostelium* are highly motile cells with dynamic, loosely bundled, short lived filopodia (Medalia et al., 2007). Given the early origins of Amoebozoa, a simple mechanism of coupling motor activation to actin dynamics at the pseudopod suggests that amoeboid filopodia formation is driven by a minimal regulatory circuit. It is tempting to speculate that VASP-dependent recruitment of an MF myosin represents an early form of cooperation between these two proteins. Perhaps as filopodia played wider roles in development and migration as multicellularity evolved, a signaling-based mechanism of MF myosin recruitment emerged with PIP3 binding to the three PH domain motif of Myo10 that replaced the first MF domain in present in other MF myosins. Thus, once Myo10 activation became dependent on PIP3, its motor activity was then used to promote VASP transport and filopodia extension. The evolution of the functional relationship between VASP and MF myosin would be interesting to explore in organisms such as *Drosophila* that make filopodia but lack Myo10 (*Drosophila* instead have a Myo22 with two MF domains like DdMyo7) and in the earliest organisms that have Myo7, Myo22 and Myo10 such as the filasterean *Capsaspora* and choanoflagellate *Salpingoeca* (Kollmar and Mühlhausen, 2017). It is possible that VASP plays a role in the recruitment of Myo10 to initiation sites or its activation in some systems, but this remains to be determined. The development of genetic tools in several evolutionarily significant organisms such as *Capsaspora* and *Salpingoeca* (Parra-Acero et al., 2018; Booth et al., 2018; Booth and King, 2020), unicellular organisms at the onset of multicellularity, should now allow for the study of the evolution of the VASP-filopodial MF myosin relationship in the targeting and activation of filopodial myosins.

## Materials and Methods

### Cell lines, cell maintenance and transformations

*Dictyostelium* control/wild-type (AX2 or AX3), *myo7* null (HTD17-1) (Tuxworth et al., 2001), *vasp* null (Han et al., 2002) and *dDia2* null (Schirenbeck et al., 2005) cells were cultured in HL5 media (Formedium). To create transgenic lines, cells were harvested, washed twice with ice cold H50 (20 mM HEPES, pH 7.0, 50 mM KCl, 10 mM NaCl, 1 mM MgSO4, 5 mM NaHCO_3_, 1 mM NaH2PO_4_ and flash spun at 10,000 X g until the rotor reached speed. Cells were resuspended at 5e7 cells/mL and 100uL of cells was combined with 10µg DNA in a 0.1 cm gap cuvette. Cells were electroporated by pulsing twice, 5 seconds apart with a Bio-Rad Gene-Pulser set to 0.85 kV, 25 µF, and 200 Ω. Cells were recovered 10 minutes on ice and plated in a 10cm dish for 24 hours before moving to selection media, either 10µg/mL G418, 35µg/mL HygromycinB or both.

Expression of the fusions proteins or null backgrounds was verified by western blotting using either anti-Myo7 (UMN87, (Tuxworth et al., 2005)), anti-GFP (Biolegend - B34) or anti-VASP (Breitsprecher et al., 2006) with anti-MyoB used as a loading control (Novak et al., 1995) (Supplemental Figure 6).

### Generation of expression plasmids

The GFP-DdMyo7 expression plasmid was created by fusing *gfp* to the 5’ end of the *myoi* gene (dictyBase DDB: G0274455; (Titus, 1999) then cloning it into the pDXA backbone with the actin-15 promotor and a NeoR cassette as described (Tuxworth et al., 2001). Plasmids used for the expression of the full length tail (aa 809 - end) (Tuxworth et al., 2001), the KKAA autoinhibition mutant (K2333A/K2336A) (Petersen et al., 2016), motor forced dimer (aa 1-1020 followed by the mouse Myo5A coiled coil region and a GCN4 leucine zipper) (Arthur et al., 2019) have been described previously. An expression clone for the full-length GFP-DdMyo7 with a C-terminal prenylation site (CAAX) was generated using Q5 mutagenesis (New England Biolabs) to add codons encoding the CTLL* prenylation motif from *Dictyostelium* RasG (UNIPROT: P15064) to the 3’ end of the *myoi* gene. A Ddyo7-mCherry expression plasmid was generated by first TA cloning a PCR product (myo42 to myi185+2) encompassing the *myoi* 3’ region of the gene (aa 475 - end) minus the stop codon using StrataClone (Agilent) (pDTi289+2). This fragment was then cloned into pDM CCherry, a modified pDM358 (Veltman et al., 2009) with the mCherry gene inserted for C-terminal fusions, generating pDTi299. The 5’ end of *myoi* was then inserted by restriction cloning to create pDTi340. A motor mutant that cannot hydrolyze MgATP, the non-hydrolyzer E386V was designed based on a characterized *Dictyostelium* Myo2 mutant (Friedman et al., 1998). The combined non-hydrolyzer + KKAA was made by standard ligation cloning to introduce the motor domain sequence from the non- hydrolyzer mutant into KKAA full length expression plasmid by restriction enzyme digest with BsiWI and BstEII. The uncoupler mutant (I426A) was based on a characterized *Dictyostelium* Myo2 mutant (Sasaki et al., 2003) was cloned by Q5 mutagenesis. Fluorescent protein fusions of DdMyo7 were made using a combination of Q5 mutagenesis, Gibson assembly and restriction enzyme cloning. A DdMyo7-Scarlet I expression plasmid, pDTi517, was generated by first cloning a full-length *myoi* gene that has a BglII site at the 5’ end, lacks its internal BglII site and also its 3’ stop codon, pDTi515+2, using a combination of Q5 mutagenesis, Gibson assembly and restriction enzyme cloning. The base Scarlet I-pDM304 expression plasmid for C-terminal fusions was generated by restriction enzyme cloning a codon-optimized synthesized Scarlet I gene (GenScript) into the extrachromosomal expression plasmid pDM304 (Veltman et al., 2009). The *myoi* gene was then cloned into mScarlet I-pDM304. Restriction cloning was used to create wild type and non-hydrolyzer mNeon DdMyo7 expression plasmids (pDTi516 and pDTi527, respectively). First, the base mNeon-pDM304 expression plasmid for N-terminal fusions was generated by restriction enzyme cloning a codon-optimized synthesized mNeon gene (GenScript) into the extrachromosomal expression plasmid pDM304 (Veltman et al., 2009). Then a full-length *myoi* gene that has a Bgl II site at the 5’ end and lacks its internal Bgl II site was generated and cloned into either pDM448 (Veltman et al., 2009) for GFP-DdMyo7 expression (pDTi492) or mNeon-pDM304 for mNeon-DdMyo7 expression (pDTi516). Restriction cloning was used to exchange the 5’ region of the gene carrying the E386V mutation into the wild type pDTi516 expression plasmid, creating pDTi527. The dDia2-CA mutant was created by first PCR-amplifying the *forH* gene (using forH4L and forH8 oligos) from GFP-dDia2 (Schirenbeck et al., 2005) then TA cloning the product using StrataClone (Agilent) to generate dDia2-SC. Mutations to generate a double R1035A, R1036A mutation were introduced dDia2- SC by Q5 mutagenesis (dDia2-CA SC). The wild type or mutant genes were restriction cloned into the extrachromosomal expression plasmid pDM449 (Veltman et al., 2009) to generate mRFPmars-dDia2 wt or CA. An inducible V-1 expression plasmid was created by first PCR-amplifying the *mtpn* gene with V-1 F and V-1 R oligos (dictyBase:DDB_G0268038) using Ax2 genomic DNA then TA cloning the product using StrataClone (Agilent) to generateV-1 SC. The V-1 insert was then restriction cloned into the extrachromosomal expression plasmid pDM334 (Veltman et al., 2009) to generate GFP-inducible V-1. The sequence of all PCR generated clones was confirmed by Sanger sequencing (GeneWiz and University of Minnesota Genomics Center).

The GFP-VASP expression plasmid was a gift from Dr. Richard Firtel (UCSD) (Han et al., 2002). The VASP tetramer and FAB, 1M mutants were not fused to GFP to avoid any steric hindrance with the fluorescent protein. The full-length VASP cDNA (dictyBase:DDB_G0289541) was cloned into the pDM344 shuttle vector (Veltman et al., 2009) and the NgoM-IV fragment from this plasmid was ligated into pDM358-mApple that has the mApple gene (Shaner et al., 2008) cloned in between the *act6* promoter and *act15* terminator of pDM358 (Veltman et al., 2009). VASP-1M was created by introducing a SmaI site and stop codon into the *vasp* gene, altering the coding sequence from 334 PSLSAPL to 334 PSLSAPG* using Q5 mutagenesis. The F-actin binding mutant (FAB K-E) was based on mutating previously identified critical F-actin binding residues (K275, R276, K278, and K280; (Schirenbeck et al., 2006) to glutamic acid (Hansen and Mullins, 2010) by Q5 mutagenesis. The sequence of all PCR generated clones was confirmed by Sanger sequencing (University of Minnesota Genomics Center). Oligonucleotides used are in the key resource table.

## Microscopy and imaging experiments

### Live-cell imaging

Microscopy of live cells was carried out as previously described (Petersen et al., 2016). Briefly, cells were adhered to glass bottom imaging dishes (CellVis, D35-10-1.5-N) and starved for 45 to 75 min in nutrient-free buffer (SB, 16.8 mM phosphate, pH 6.4), and then imaged at 1 to 4 Hz on a spinning disk confocal (3i Marianas or Zeiss AxioObserver Z.1) with a 63 X (1.4NA objective, 0.212 micron pixel size). The sample temperature was maintained at 19 - 21°C. Samples were illuminated with 50mW lasers (488nm or 561nm), and a Yokogawa CSU-X1 M1 spinning disk, and captured with an Evolve EMCCD camera. 4 - 6 Z sections of 0.28 - 0.5 microns were taken with a 50 - 250ms exposure with 10-40% laser power. Cells were imaged for 10 seconds – 10 minutes or longer depending on experiment. Cells are plated at a density of 5×10^5^ per mL. Ten fields of view were collected from each imaging dish, with 2 - 20 cells per field of view. All data sets represent cells from at least three independent experiments and two independently transformed cell lines.

### Drug treatments

Cells were washed free of media, adhered to glass bottom dishes and starved in nutrient-free buffer for 40 minutes. The buffer was replaced by buffer supplemented with the noted concentration of Jasplakinolide (diluted to 0.5% DMSO), cytochalasinA, latrunculin A, CK666, nocodazole, Ly294002, wortmannin or just DMSO alone. Jasplakinolide treatment was for 5 - 8 minutes, cells were incubated with all other compounds for 15 - 20 minutes, prior to imaging for 10 - 30 minutes. Additional drug concentration data are in Table 1. Cells expressing mApple-DdMyo7 and inducible GFP-V1 were induced overnight with 10µg/mL doxycycline (Sigma) to turn on expression of V-1 prior to imaging as above. Cells were treated with 0.25 µg/mL FM4-64 (Invitrogen) for 2 - 5 minutes to image filopodia with a membrane marker.

### LatrunculinA-induced actin waves

Cells expressing GFP-DdMyo7 and the actin reporter RFP- LimEΔcoil (Gerisch et al., 2004) were induced to generate travelling actin waves using a modified protocol (Gerisch et al., 2004). Cells rinsed with 16mM phosphate buffer pH 6.4 were seeded on glass bottom dishes (Celvis) at 5×10^5^ cell/mL, incubated in phosphate buffer for 30 minutes and then supplemented with 5µM latrunculinA for 20 minutes. The solution was diluted to 0.5µM latA and cells incubated for an additional 30 minutes, then imaged for up to 2 hours. Images were captured in 5 - 10 0.3 µm Z sections by spinning disk confocal microscopy (see above) every 5 seconds to make 10 - 30 minute movies.

### Actin intensity linescans

Cells were seeded as above and fixed using picric acid (Humbel and Biegelmann, 1992). Cells were incubated with 568- or 647- Alexa phalloidin (Invitrogen) for 45 minutes, rinsed with PBS-glycine, then water and mounted using prolong-diamond (Molecular Probes). Slides were imaged using a Nikon Widefield Eclipse NiE microscope with a 40X/1.3 NA objective, CoolSNAP CCD camera and Sola light source. Maximum Z projections (5 - 10 0.3 µm) were analyzed by manually drawing a linescan perpendicular to the long axis and compiled with the FIJI/ImageJ macro and RStudio scripts (Zonderland et al., 2019).

## Data Analysis

### Protein alignments

Myosin motor domain sequences were aligned using T-coffee algorithm “Expresso” (Notredame et al., 2000). These included Uniprot sequence entries: P54697, Q9U1M8, P08799,P19524, K4JEU1, Q9V3Z6, Q9HD67,Q13402, Q6PIF6. Diaphanous related formins were aligned using Clustal Omega, using Uniprot entries: Q54N00, O70566, O60879, P41832, P48608 and the DAD domain basic residue highlighted is based on (Wallar et al., 2006; Lammers et al., 2005). VASP mutations were made by first creating a structural alignment with DdVASP (Uniprot: Q5TJ65) and Human VASP (Uniprot: P50552) in T-coffee algorithm “Expresso”.

### Image Analysis

Cortex to cell ratio, cortical asymmetry and filopodia per cell were quantified using a custom FIJI plugin “Seven” ((Petersen et al., 2016), code available on github linked to titus.umn.edu). Cells not expressing transgenic proteins were excluded from the analysis. Analysis was done on maximum intensity projections. First, the image is thresholded to mask the cell. This mask excludes the nucleus, which is devoid of DdMyo7 signal, and filopodia tips extended from the cell body. The cortical band (0.8µm) intensity, standard deviation of intensity in the cortical band was measured for each cell. Next, a radial tip search identifies filopodia and registers them to each cell to count the number of filopodia per cell. Extending pseudopodia and actin correlation were done in FIJI by reslicing through extending pseudopodia and making plot profiles of the edge of the cell. Intensity profiles were normalized between 0-1, and multiple cells were averaged by fitting a restricted cubic spline.

### Statistical analysis

Statistics were calculated in Prism 8 (GraphPad). One-way ANOVA analysis with post hoc Tukey test or Dunnett’s multiple comparison to wild-type control was used to compare groups; Student’s t test was used when only comparing 2 datasets. Statistical tests were calculated on full datasets, experimental means are shown on graphs to demonstrate experimental variability. Automated analyses data points deemed definite outliers (0.1%) by Rout method were excluded for cortex:cytoplasm ratio. Error bars are SEM, unless noted. Significance differences are in comparison to control (DdMyo7), unless noted. Tables have filopodia number as average number of filopodia in cells with at least one filopod. SEM is standard error of the mean. Capitalized ‘N’ indicates number of experiments lowercase n is number of cells.

### Cytoskeleton Association

Log phase cells expressing GFP-DdMyo7 were grown on bacteriological plastic plates (150 mm) were rinsed twice in PB then resuspended, counted and 5 x10^7^ cells collected by centrifugation (Beckman J6, 200 x g). The pellet was resuspended in 1 ml lysis buffer (100 mM Tris, pH 8.1, 5 mM MgCl_2_, 5 mM EDTA and 2.5 mM EGTA), washed once then lysed with Lysis Buffer + 1% Tx-100, 1 mM TLCK (Sigma), 1 mM TPCK (Sigma), 1X HALT protease inhibitor cocktail (Pierce ThermoFisher) at room temperature. The 0.5 ml sample was spun immediately at 20,000 x g, 4°C for 20 min. The supernatant was collected and pellet resuspended and homogenized in 0.25 ml of lysis buffer. An equal volume of ULSB gel sample buffer was added to aliquots from the sup and pellet and the samples run on a 4 - 15% gradient TGX SDS PAGE gel (BioRad) then transferred to nitrocellulose (Licor). The blot was probed for the presence of DdMyo7 and actin followed by fluorescent secondary antibodies (LiCor) and then imaged with the LiCor Odyssey. Quantification of the DdMyo7 band was performed using Image Studio Lite (Licor).

## Key Resources

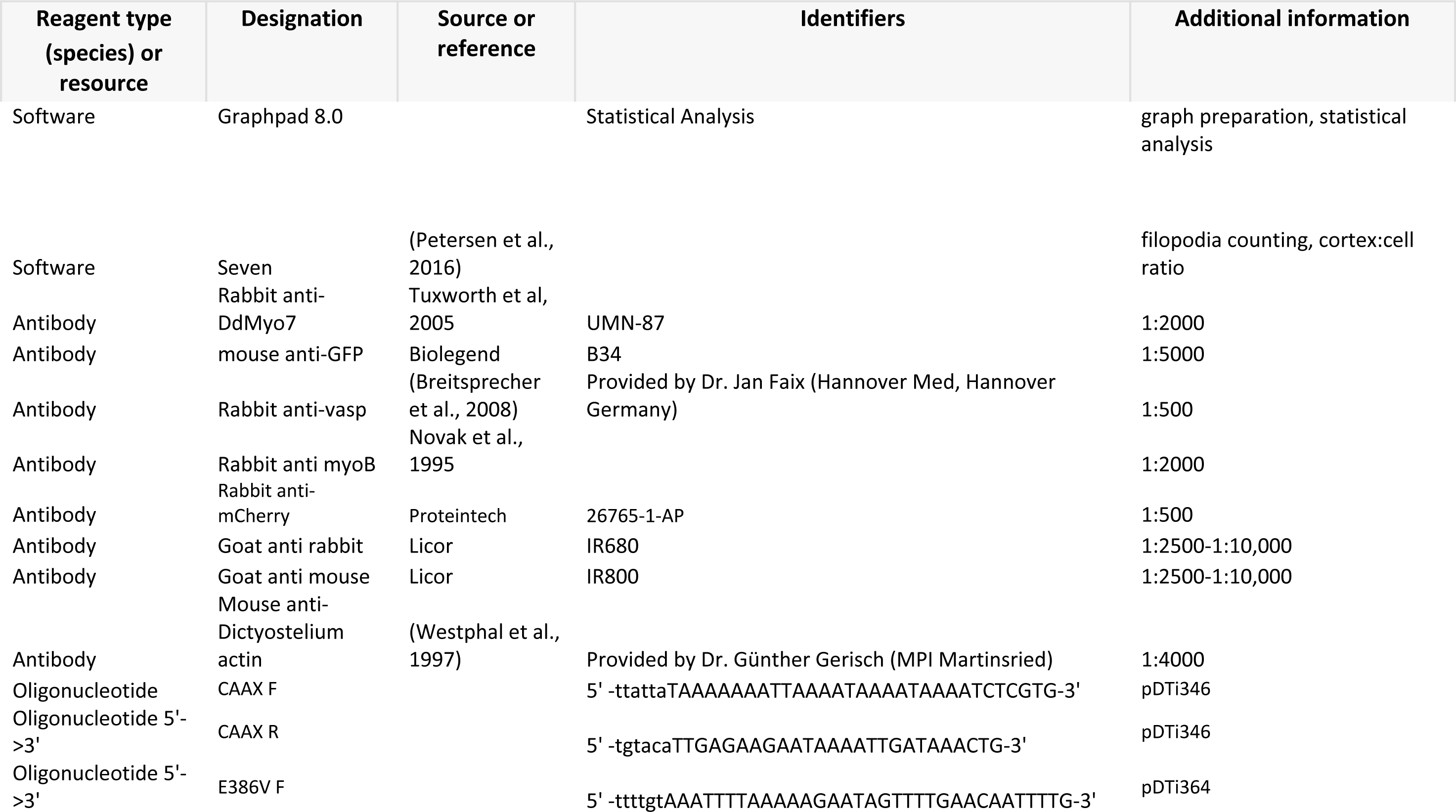

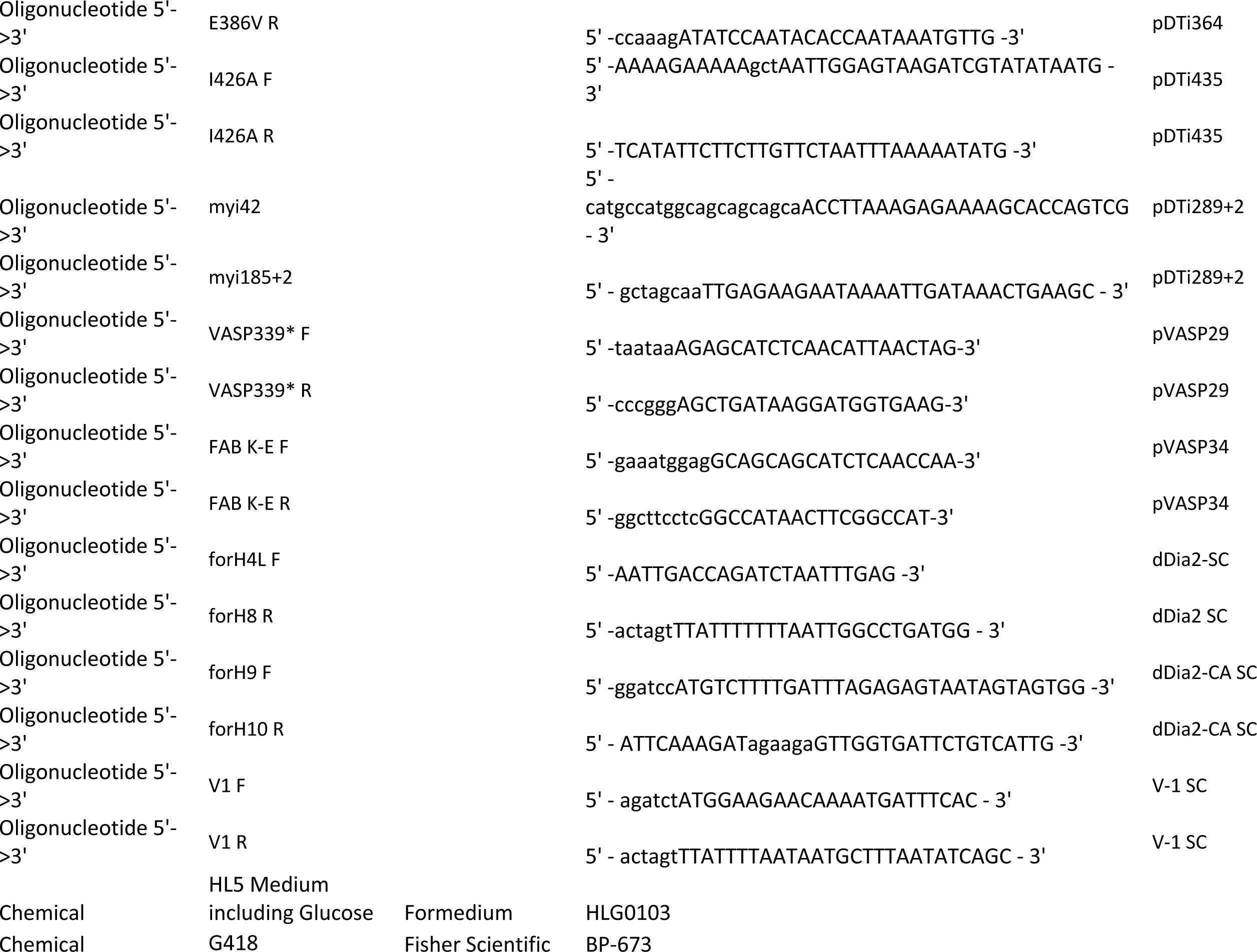

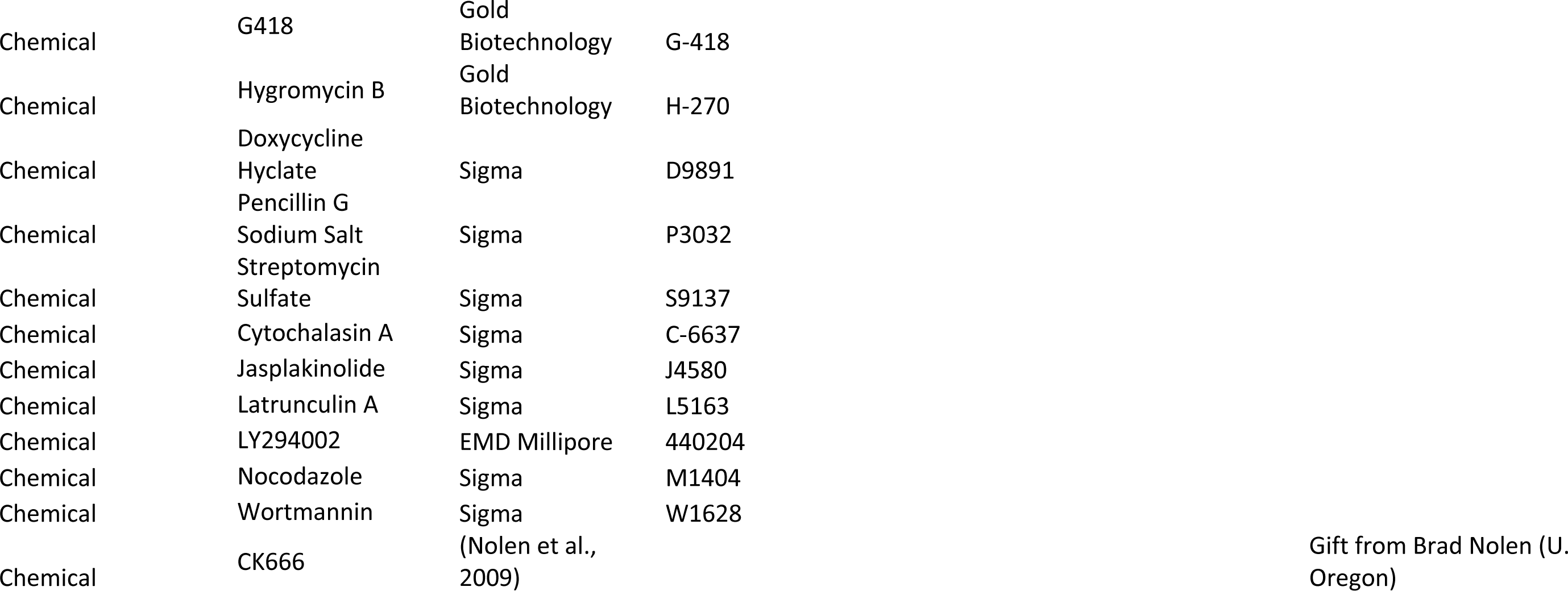

synthesized DNA mScarlet I

actagtggtggttcaggaGTTTCAAAAGGTGAAGCCGTTATTAAAGAATTTATGAGATTCAAGGTTCACATGGAAGGAAGTATGAACGGTCATGAATTTGAGATTGAAGGAGAAGGTGAAGGTAGACCATATGAAGGCACCCAAACAGCTAAATTAAAAGTAACTAAAGGTGGTCCATTACCATTTAGTTGGGATATTTTATCTCCACAATTTATGTATGGTTCACGTGCTTTCAttAAACATCCAGCAGATATTCCAGATTATTATAAACAATCATTTCCAGAAGGTTTTAAATGGGAACGTGTCATGAACTTTGAAGATGGTGGAGCAGTTACAGTCACACAAGATACCTCATTAGAAGATGGTACATTAATATATAAAGTTAAATTACGTGGTACTAATTTTCCACCAGACGGTCCAGTAATGCAAAAAAAAACAATGGGCTGGGAAGCTAGTACAGAACGTTTATATCCTGAAGATGGTGTCCTTAAAGGCGATATAAAAATGGCCTTGAGATTAAAGGATGGTGGTAGGTATTTAGCAGATTTCAAAACCACTTATAAAGCAAAAAAACCAGTTCAAATGCCAGGTGCATATAATGTTGATAGAAAACTTGATATTACCAGTCATAATGAAGATTACACAGTTGTCGAACAATACGAACGTTCTGAAGGTCGTCATAGCACTGGTGGTATGGATGAATTATACAAATAAgctagc

synthesized DNA mNeon

ggatccATGGTGAGTAAAGGTGAAGAAGATAATATGGCATCGTTACCAGCTACACATGAGTTACATATATTCGGTAGCATTAATGGTGTTGATTTTGATATGGTGGGACAAGGTACCGGTAATCCTAATGATGGTTACGAAGAACTAAATTTAAAATCGACTAAAGGTGACTTACAATTTTCTCCATGGATTTTAGTGCCACATATAGGGTATGGTTTTCATCAATACTTACCATATCCAGATGGTATGTCACCATTTCAAGCTGCAATGGTTGATGGATCAGGTTATCAAGTTCATAGAACAATGCAATTTGAAGATGGTGCTTCATTAACTGTTAATTATAGATACACATATGAAGGCTCACATATTAAAGGTGAAGCTCAAGTTAAAGGTACTGGTTTCCCAGCCGATGGCCCAGTTATGACAAATAGTTTAACAGCAGCAGATTGGTGTAGATCCAAAAAAACTTATCCAAATGATAAAACAATTATTTCAACTTTTAAATGGTCATATACAACCGGTAATGGTAAACGTTATCGTTCAACAGCCCGTACAACATATACTTTTGCTAAACCAATGGCAGCTAATTATTTAAAAAATCAACCAATGTATGTTTTTCGTAAAACAGAGTTAAAACATTCAAAAACAGAACTTAATTTTAAAGAATGGCAAAAAGCATTTACAGACGTTATGGGTATGGATGAACTTTATAAGAgatct

## Acknowledgements

We thank Livia Songster, Annika Schroder and Casey Eddington for many helpful discussions and assistance with experiments and analysis. We also thank Dr. Karl Petersen for the initial characterization of the motor mutations and for support with SEVEN. We are grateful to Professor Robert Insall (Beatson) for stimulating discussions, Dr. John Cooper (Washington U), Dr. Volodya Gelfand (Northwestern U), Dr. Lil Fritz-Laylin and members of the Fritz-Laylin lab (UMass Amherst) for helpful comments and critical reading of the manuscript. Thanks for Dr. Jan Faix (U Hannover) for providing the dDia2 nulls and VASP antibody and Dr Brad Nolen (U. Oregon) for supplying the CK666. Our thanks also to the University of Minnesota Imaging Center for additional imaging support. This work was supported by the CNRS, ANR-17-CE11- 0029-01, and Ligue Contre le Cancer RS16 grants (A.H.). The A.H. team is part of the LabexCelTisPhyBio:11-LBX-0038, which is part of the Initiatives of Excellence of Université Paris Sciences et Lettres (ANR-10-IDEX-0001-02PSL) and the NIH National Institute of General Medical Sciences (F31GM128325 to A.L.A. and R01GM122917 to M.A.T.).

---

**Supplemental Figure 1.**
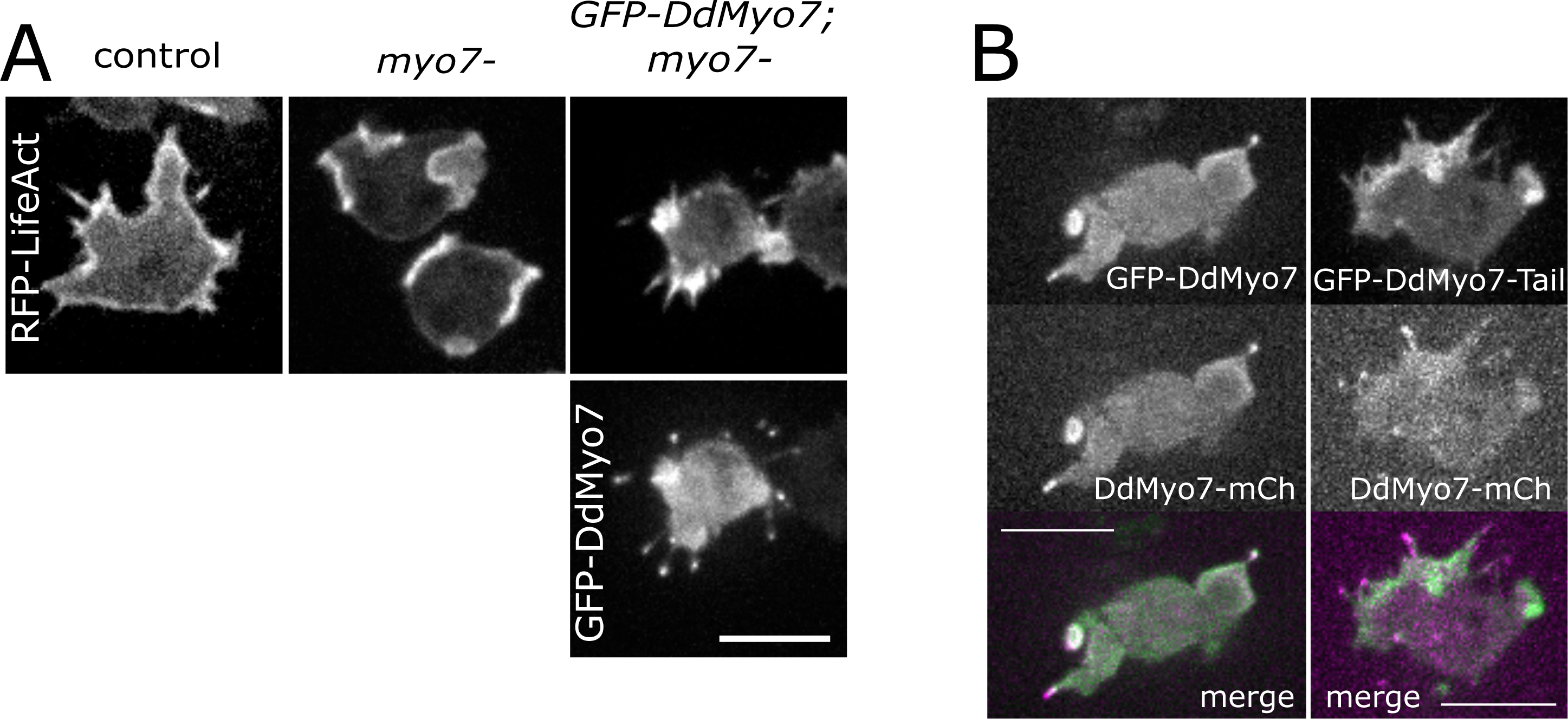
**A.** Micrographs of control (AX2), *myo7* null or *myo7* null with GFP- DdMyo7 rescue construct expressing RFP-LifeAct (actin, top) and by GFP-DdMyo7 (bottom). **B.** Micrographs of cells co-expressing either GFP-DdMyo7 and DdMyo7-mCherry or GFP- DdMyo7-tail and DdMyo7-mCherry. Scale bars: 10µm.

**Supplemental Figure 2.**
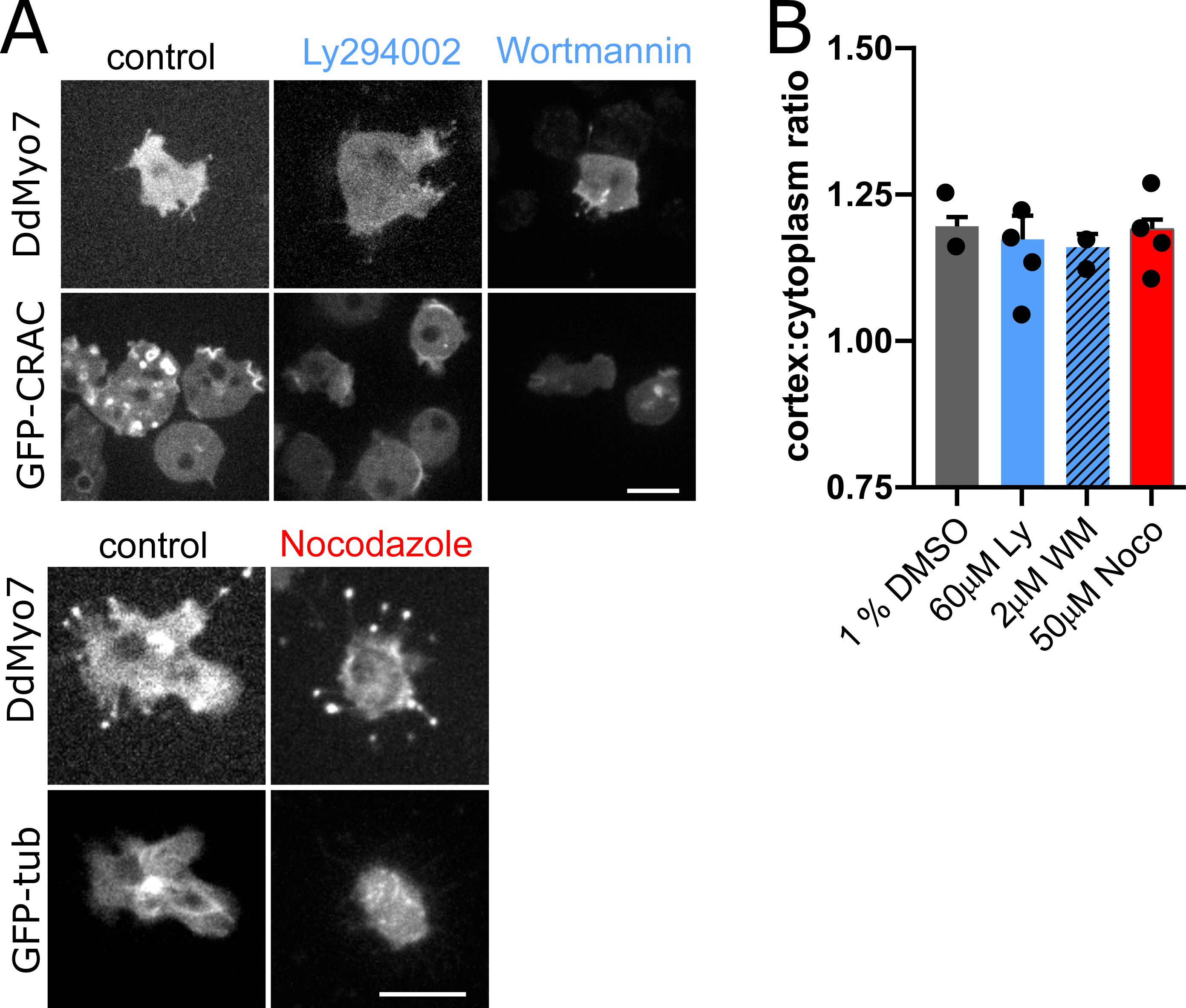
**A.** Micrographs of cells expressing DdMyo7 and GFP-CRAC (top) or GFP-tubulin (bottom) under noted drug condition. **B.** Quantification of additional pharmacological compounds on DdMyo7 cortical recruitment, cells were incubated with Ly294002 (Ly), Wortmannin (WM) or nocodazole (Noco). Circles are experimental means, one-way ANOVA indicates no data are significantly different from control. Scale bar is 10µm.

**Supplemental Figure 3.**
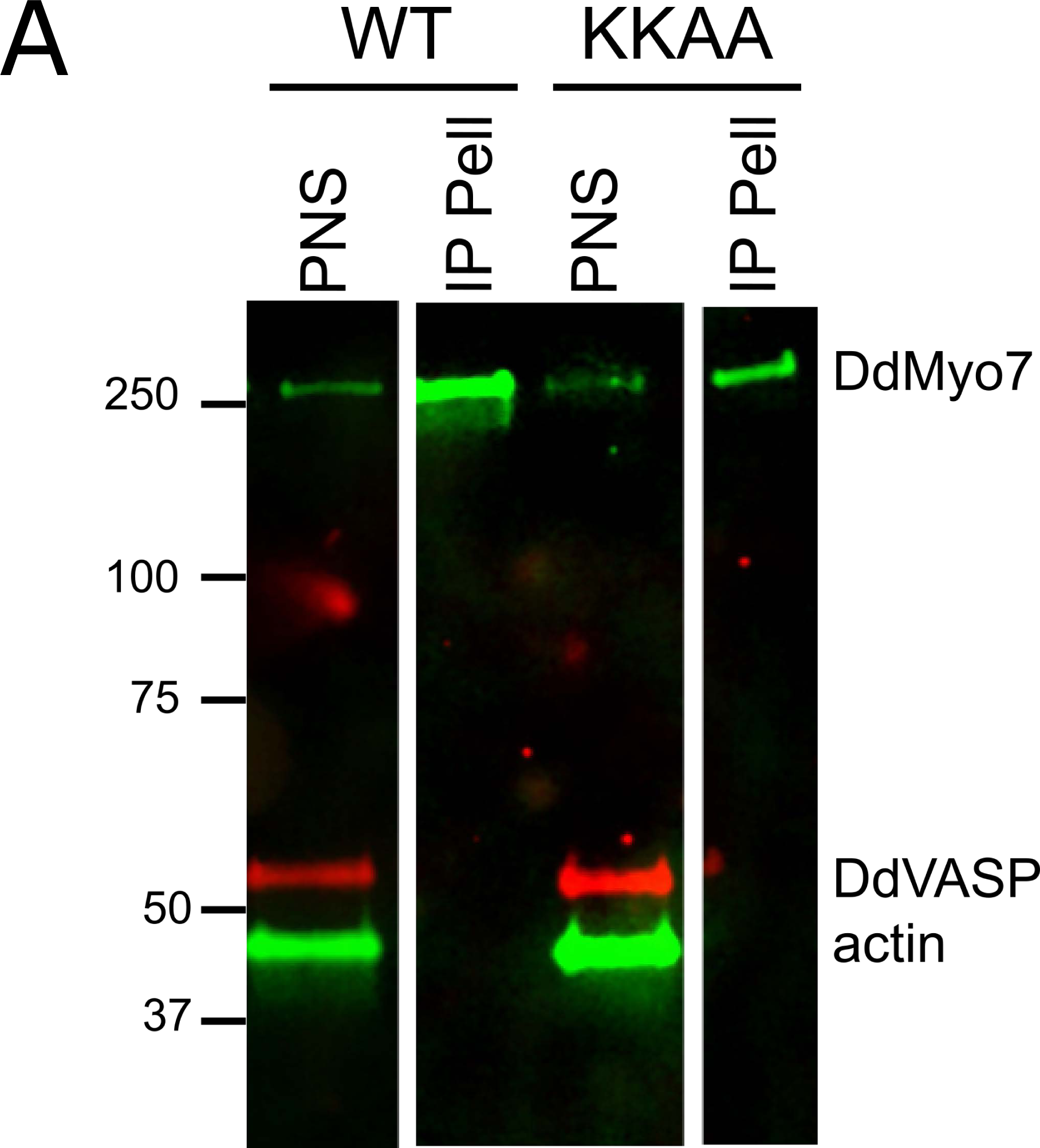
Immunoprecipitation of DdMyo7. **A.** GFP-DdMyo7 or GFP- DdMyo7-KKAA (see Fig 5) were immunoprecipitated from a clarified lysate (post nuclear spin sup; PNS) using anti-GFP beads. The PNS and immunoprecipitate pellet (IP Pell) were probed by western blot for DdMyo7, actin and DdVASP.

**Supplemental Figure 4.**
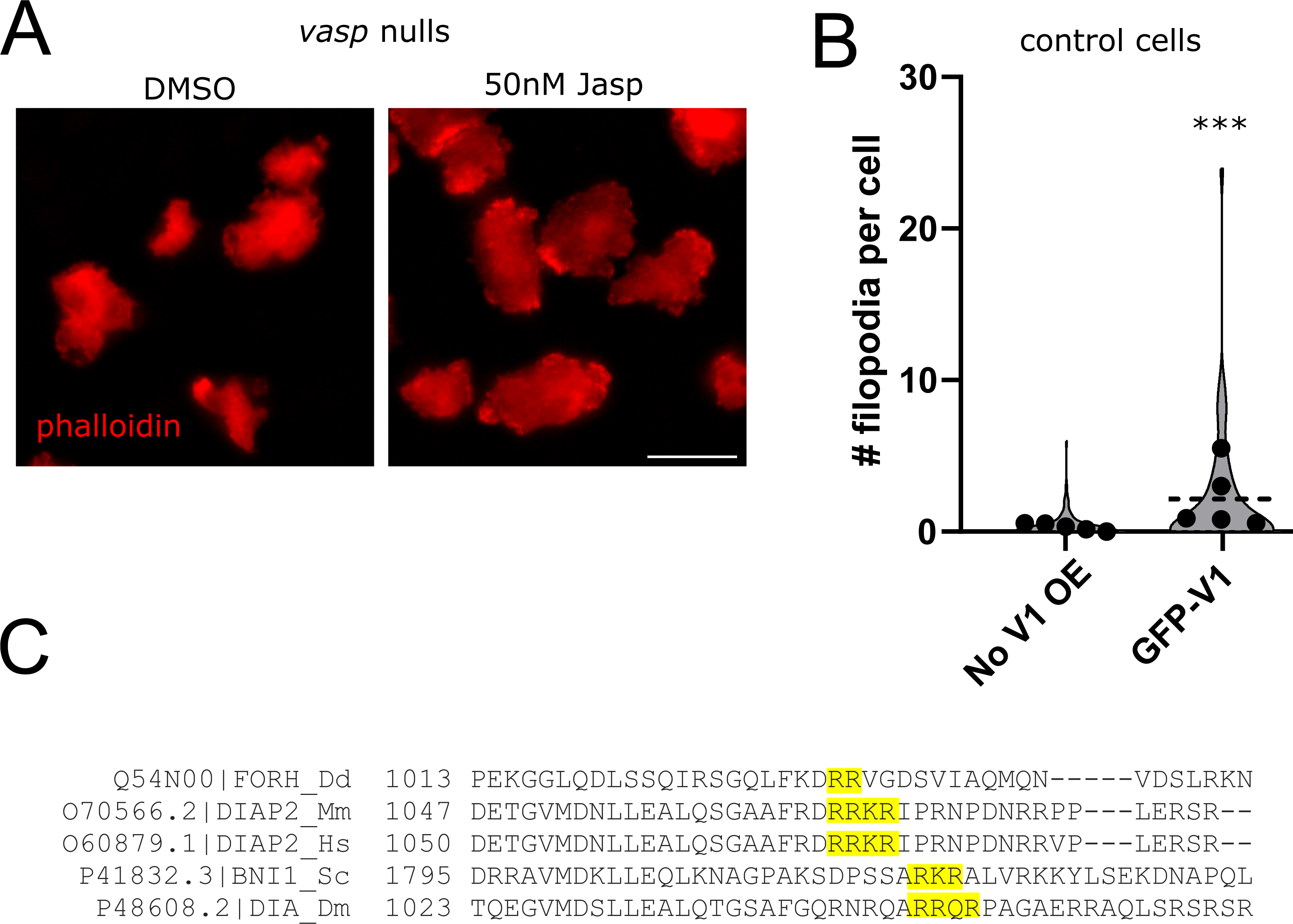
**A.** Phalloidin staining of *vasp* null cells treated with either DMSO or 50nM Jasplakinolide (Jasp). **B.** Quantification of induction of filopodia formation by control cells (no V1 OE) or cells that overexpress GFP-V1. Students t-test, ***p<0.001. **C.** Clustal Omega alignment of diaphanous related formins, conserved basic residues in the Dia autoregulatory domain (DAD) highlighted in yellow.

**Supplemental Figure 5.**
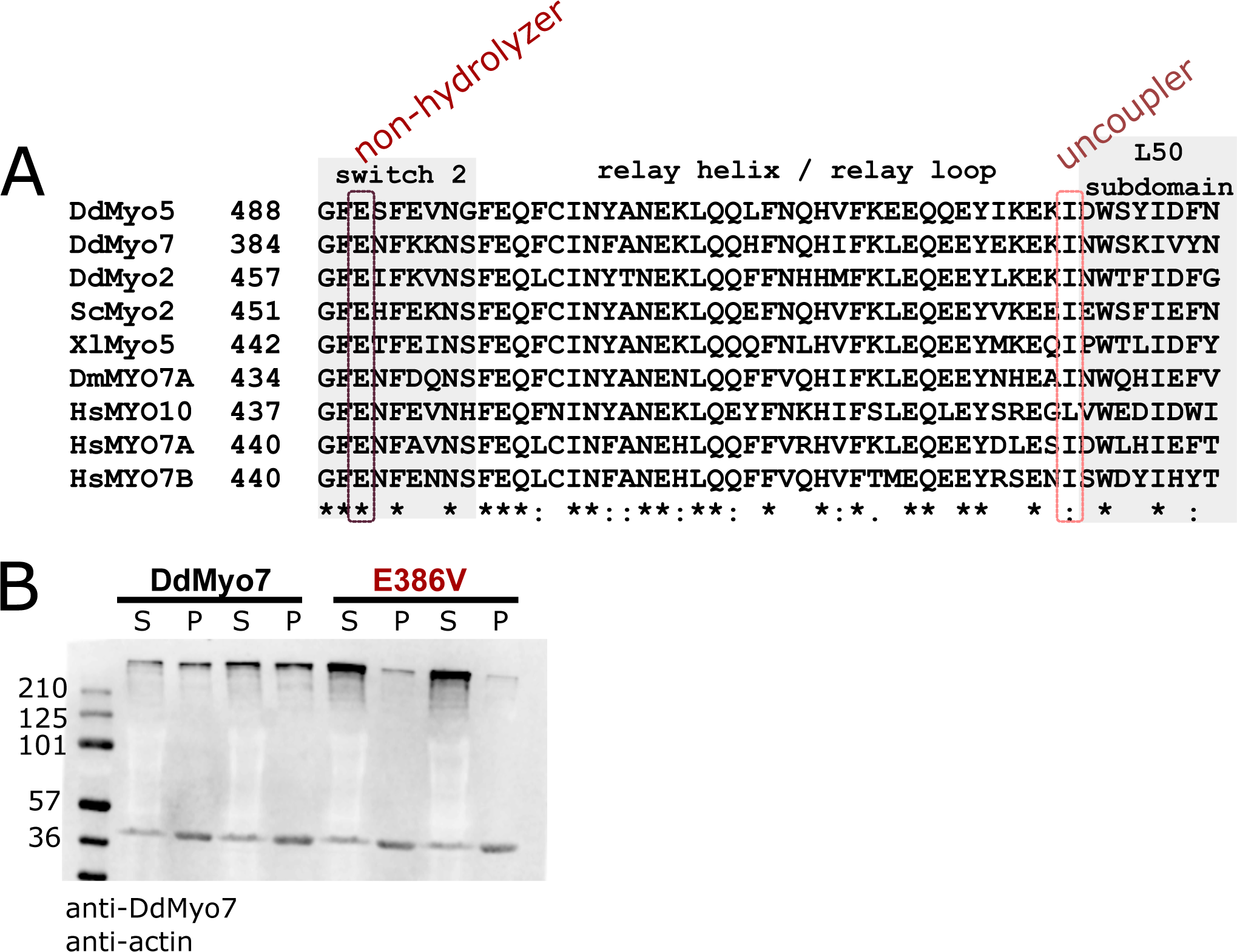
Conservation of the DdMyo7 motor domain. **A.** M-coffee sequence alignment (Wallace et al., 2006) of the relay helix region of different myosin motor domains, left column number is amino acid position. Switch 2 and L50 subdomain are shaded, circled columns indicate the highly conserved glutamic acid in switch 2 (non-hydrolyzer, DdMyo7 E386V) and hydrophobic residue in relay loop (uncoupler, DdMyo7 I426A). Symbols below indicate degree of conservation between sequences:‘*’ identical, ‘:’ strongly similar, ‘.’ weakly similar. **B.** Western blot analysis of two cytoskeleton prep supernatants (S) and pellets (P) from *myo7* null cells expressing either wildtype or the E386V mutant. Band at 270kDa is DdMyo7, band at 42kDa is actin.

**Supplemental Figure 6.**
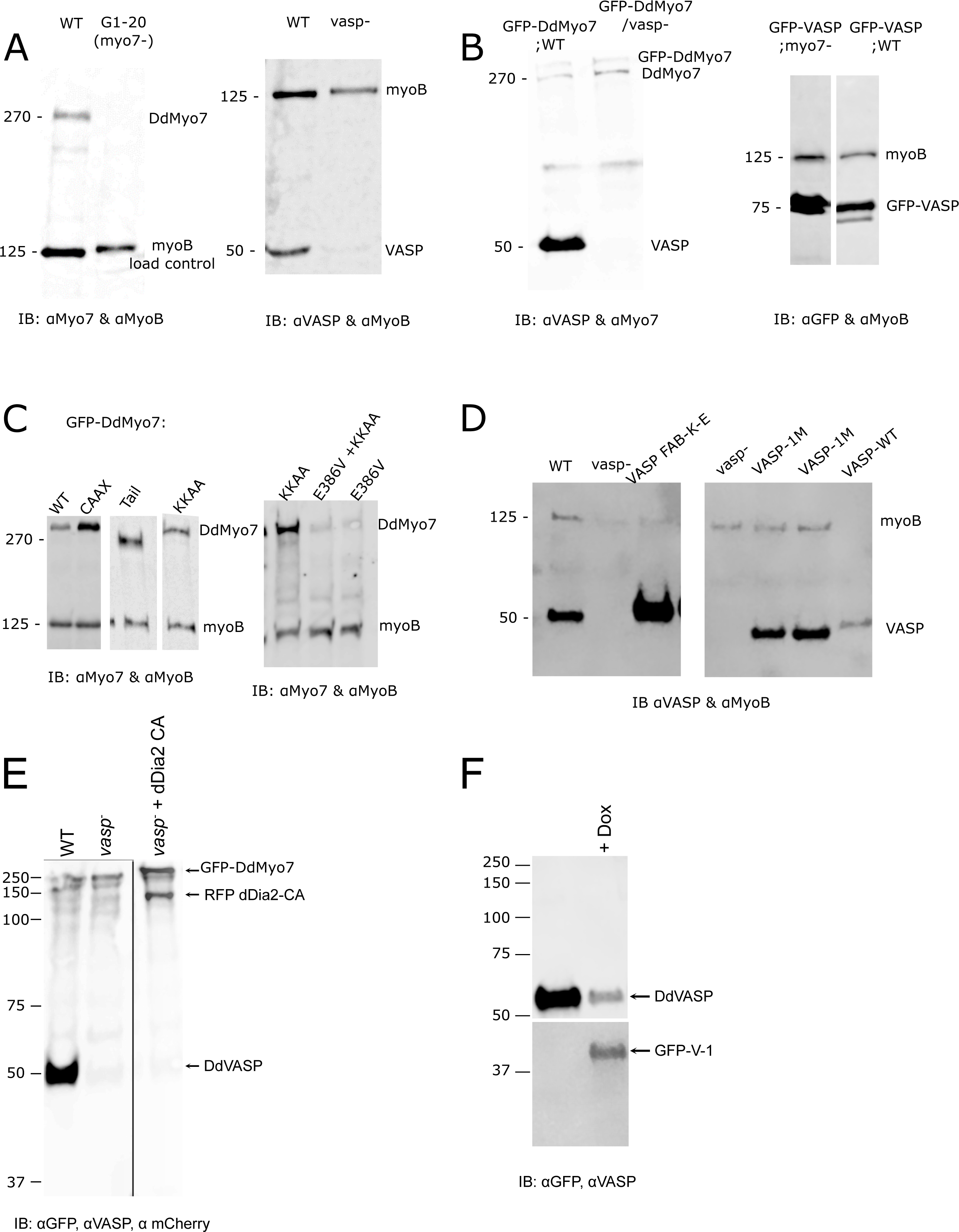
Whole cell lysates from each line were analyzed for expression of endogenous proteins or expressed fusion proteins. The blot was also probed for the 125 kD MyoB heavy chain, the loading control. Antibodies used to probe each set of blots are indicated below and the molecular weights in kD marked on the side. **A.** Control wild type (WT, Ax2), *myo7* null or *vasp* null cell lines. Note that DdVASP runs at ∼ 50 kD, higher than its calculated molecular weight of ∼ 40 kD. **B.** GFP-DdMyo7 expression in control (WT, Ax3) and *vasp* null cells, and GFP-VASP in control wild type (Ax2) and *myo7* null cells. **C.** Expression of wild type or mutant GFP-DdMyo7 in *myo7* null cells. Note that GFP-DdVASP runs at ∼ 75 kD, higher than its calculated molecular weight of ∼ 65 kD **D.** Expression of wild type VASP and VASP mutants (not fused to a fluorescent protein) in *vasp* null cells. **E.** Wild type, *vasp* null and *vasp* null cell line overexpressing GFP-dDia2 CA. **F.** Western blot of GFP-V1 induced (+Dox) in control (Ax3) cells. Average number of filopodia per cells from cells with at least one filopodia. Cortex:cytoplasm ratio is intensity ratio of a 0.8µm band around the periphery compared to the cytoplasm. N is number of experiments, n is number of cells. SEM is standard error of the mean. GFP-VASP and GFP-DdMyo7-Tail fail to efficiently target to filopodia tip and were not counted in this analysis.

